# Novel pESI-encoded autotransporter adhesin PeaP of epidemic *Salmonella* strains mediates adhesion, atypical biofilm formation, and poultry colonization

**DOI:** 10.64898/2026.05.20.725250

**Authors:** Laura Elpers, Benita Schmeißer, Pascal Felgner, Marie Köttermann, Victoria Drauch, Claudia Hess, Nikola Köpp, Lena Lüken, Michael Hess, Ohad Gal-Mor, Michael Hensel

## Abstract

*Salmonella enterica* serovar Infantis (SIN) has rapidly become the dominant serovar in poultry worldwide, a success largely linked to the acquisition of the 285 kb megaplasmid pESI. While pESI-encoded antibiotic-resistance and iron-uptake systems are well characterized, pESI-mediated adhesion mechanisms remain poorly understood. Here we identify a novel pESI-encoded monomeric autotransporter adhesin, designated PeaP (pESI-encoded autotransporter protein), and demonstrate its pivotal role in atypical biofilm formation, interference with motility, and colonization of the chicken host. Biofilm assays revealed that pESI-harboring strain SIN 119944 forms robust biofilms at 37 °C and 42 °C, temperatures at which CsgD-dependent biofilm formation is negligible. Deletion of *csgD* did not impair this phenotype, whereas deletion of *peaP* abolished high-temperature biofilm development and restored motility to wild-type levels. Proteomic profiling of sessile versus planktonic cells highlighted PeaP as the most abundant pESI-derived protein in the biofilm fraction. AlphaFold-based modelling and negative-stain transmission electron microscopy showed that PeaP comprises a C-terminal β-barrel and a 1,500 aa passenger domain with three tandem repeats, projecting filamentous appendages ∼37-nm from the outer membrane. Antibody blockade of PeaP reduced surface adhesion >6-fold, confirming its adhesive function. In an infection model of 2 day-old chicken, the *peaP* mutant displayed significantly lower colonization, indicating PeaP-mediated adhesion *in vivo*. Collectively, pESI-positive SIN deploys PeaP for CsgD-independent, temperature-tolerant biofilm formation and enhanced gastrointestinal colonization, providing a mechanistic basis for the epidemic spread of this multidrug-resistant pathogen in poultry.

## Introduction

*Salmonella enterica* is a zoonotic gastrointestinal pathogen with remarkable capacity to adapt to different hosts, to cause diseases of varying severity, and to escape immune defenses of host organisms. Additionally, *S. enterica* possess a complex array of virulence factors that enable invasion of non-phagocytic mammalian cells, and subsequently achieve an intracellular lifestyle within a membrane-bound compartment in infected host cells (Han et al., 2024). Prior to these dedicated interactions with host cells, *S. enterica* must first establish contact and adhesion to their surfaces. These requirements are met by a set of diverse adhesion factors, known as adhesiome. For *S. enterica,* the adhesiome is surprisingly complex, with up to 12 fimbrial adhesins of the family of chaperone/usher adhesins, the fimbrial adhesin Curli belonging to the secretion/precipitation class, two non-fimbrial adhesins secreted by cognate type I secretion systems, and a large number of monomeric or trimeric outer membrane proteins with adhesive properties (Hansmeier et al., 2017; Nuccio & Baumler, 2007; Wagner & Hensel, 2011). The composition of the adhesiome varies between serovars of *S. enterica* and even between strains within a specific serovar, and the underlying mechanism of this variation is the presence of adhesion genes on accessory genome elements, such as pathogenicity islands and plasmids (Sia et al., 2025).

In addition to conferring adhesion of single bacterial cells to biotic or abiotic surfaces, adhesins also contribute to multicellular properties of bacteria such as biofilm formation and persistence in the environment. Biofilm formation is a ubiquitous microbial property, and initiates with adhesion of single planktonic cells to a suitable surface, followed by a switch to sessile lifestyle, and then leads to proliferation of multicellular communities that jointly produce a biofilm matrix (Sauer et al., 2022). The biofilm matrix is commonly composed to proteinaceous components such as Curli, and polysaccharides such as cellulose. The mechanism of biofilm formation by *S. enterica* and *E. coli* have been investigated in great detail, revealing that synthesis of matrix components Curli and cellulose is controlled by master regulator CsgD (Brombacher et al., 2006). This pathway is conserved between *S. enterica* and *E. coli*, and biofilm formation is most pronounced at temperatures of 30 °C and below, and under conditions of nutritional limitation. Biofilm formation is also an important virulence-associated phenotype because biofilms within host organisms enable bacterial persistence, protection from humoral immunity, and biofilms act as source of pathogen reseeding and dissemination (Espinoza-Erazo et al., 2025).

The prevalence of *S. enterica* in humans and in livestock is highly dynamic, and strains of *S. enterica* that successfully colonize livestock can be transmitted to humans more easily via the food chain. Recently emerging strains of *S. enterica* serovar Infantis (SIN) harbor a megaplasmid designated as ‘plasmid of Emerging *Salmonella* Infantis’ in short pESI (Aviv et al., 2014; Gal-Mor et al., 2010). These isolates were first described in Israel and associated with dominant presence in poultry as well as human infections (Aviv et al., 2016). Since then, pESI-positive SIN strains spread world-wide (Aviv et al., 2016; Cohen et al., 2022a), and a global populational WGS analysis confirmed the presence of pESI in 71% of poultry isolates and already 32% of human isolates (Mattock et al., 2024). With a size of 285 kb, pESI is a rather complex genetic unit that is self-transmissible and maintained by at least three toxin/antitoxin systems. pESI confers resistance to the antibiotics tetracycline, streptomycin, spectinomycin, sulfonamides, resistance to disinfectants of the quaternary ammonium compound family, and heavy metal mercury (Aviv et al., 2014; Cohen et al., 2020). Interestingly, SIN managed to occupy the niche of broiler production systems, leading to the most common serovar in broilers and broiler meat in the European Union (EFSA, 2025). Furthermore, a strong association between the presence of pESI and epidemic success of *Salmonella* clones has been observed (Alba et al., 2020; Cohen et al., 2022b; Guzinski et al., 2023; Mattock et al., 2024). In order to understand the epidemic success of SIN [pESI], we set out to identify pESI-encoded functions mediating adhesion. Two fimbrial adhesins encoded by genes in pESI have recently been described (Aviv et al., 2017), and increased biofilm formation was reported for SIN [pESI] (Aviv et al., 2014). Our current work identifies a novel pESI-encoded autotransporter adhesin termed PeaP, and describes its adhesive and structural properties. We furthermore demonstrate the role of PeaP in CsgD-independent, atypical biofilm formation, and its importance during colonization of the chicken gastrointestinal tract.

## Results

### Salmonella Infantis harboring megaplasmid pESI exhibit atypical biofilm formation

In search for factors that contribute to the remarkable global spread of pESI-positive SIN isolates (SIN [pESI]), we investigated the role of pESI in biofilm formation. A previous study on the characterization of SIN [pESI] reported increased biofilm formation by SIN [pESI] isolates compared to pre-epidemic isolates (Aviv et al., 2014). The previous study determined biofilm formation on cholesterol-coated plastic surfaces after 48 h at 37 °C. To further corroborate this observation, we quantified biofilm formation on abiotic surfaces under various environmental conditions. Biofilm formation by *S. enterica* serovar Typhimurium (STM) is known to be controlled by the master regulator CsgD, involves Curli fimbriae and extracellular polysaccharide (EPS) cellulose as main components of the biofilm matrix, and highest levels of biofilm formation were observed at 30 °C or lower in nutrient-deprived media. (Gerstel & Römling, 2003; Liu et al., 2014). In biofilm assays using CFA medium and non-treated polystyrene multi-well plates, we determined CsgD-dependent biofilms of STM at 30 °C, but not at 37 °C or 42 °C (**Figure 1**). The mass of STM biofilm increased during an incubation period of up to 144 h. Similar biofilm formation at 30 °C was determined for SIN 119944 (emerging isolate harboring pESI) and SIN 335-3 (pre-emerging isolate lacking pESI), but for SIN 119944, high levels of biofilm were already present after incubation for 24 h at 30 °C (**Figure 1**). At higher temperatures of 37 °C or 42 °C, STM and SIN 335-3 did not form biofilms under these assay conditions, whereas SIN 119944 maintained biofilm formation reaching maximum level already after 24 h of incubation.

**Figure 1.**
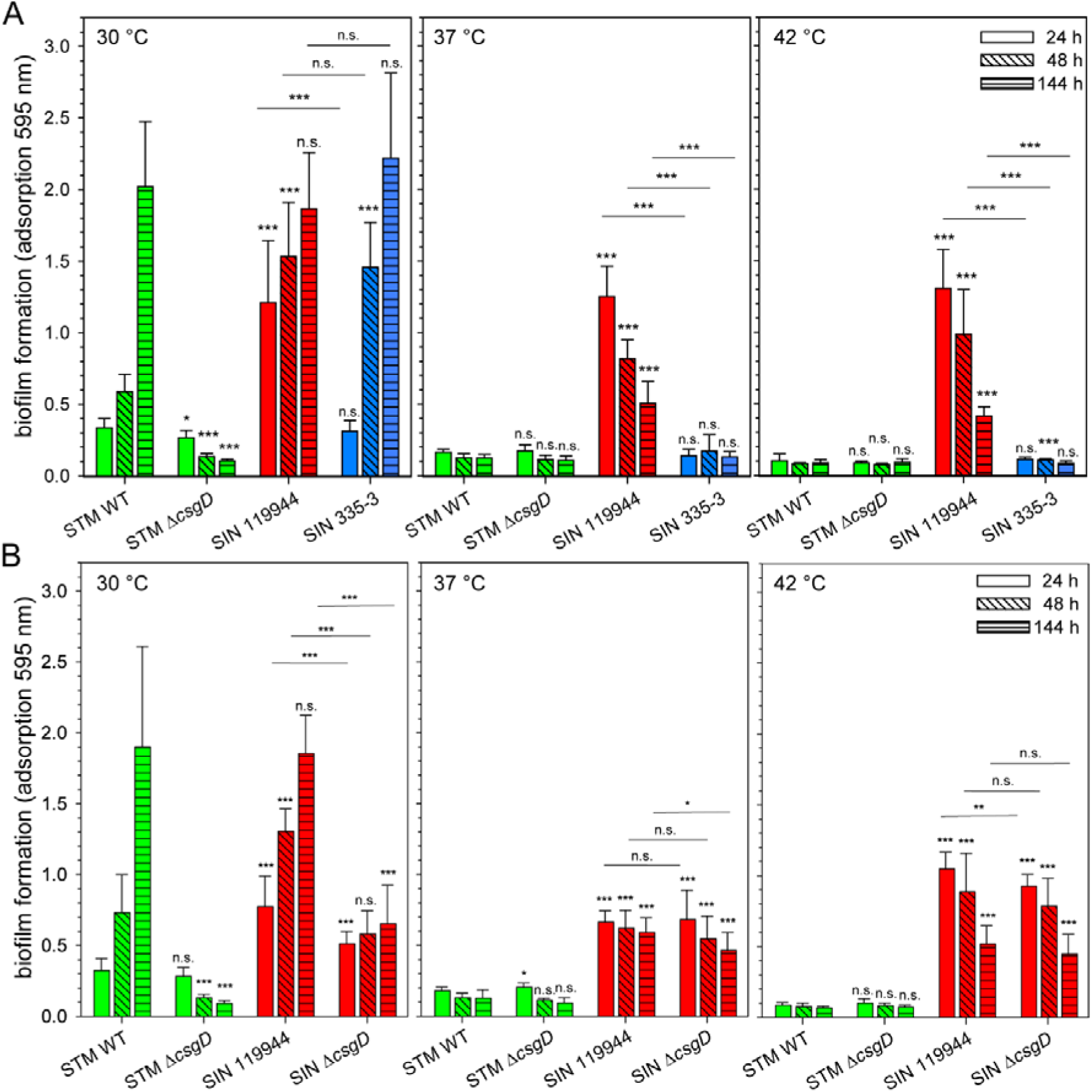
Emergent SIN 119944 shows increased biofilm formation at higher temperatures that is independent of master regulator CsgD. **A**) STM and SIN strains were grown o/n and subcultured for 3.5 h in CFA medium at 37 °C with aeration. Suspensions containing ca. 1 x 10^8^ bacteria/ml in CFA medium were dispensed in wells of 96-well plates in triplicate and incubated statically in a humidified chamber at 30 °C, 37 °C, or 42 °C as indicated. As control, wells with sterile CFA medium were used. After incubation for 24 h, 48 h, or 144 h, medium was aspirated, wells washed thrice, and biofilm formation was quantified by crystal violet staining and determination of absorbance at 595 nm using a multi-well plate reader (Chameleon X). Means and standard deviations of at least three biological replicates are shown. **B**) Biofilm formation of STM, SIN 119944, and isogenic mutant strains deficient in biofilm master regulator *csgD* were analysed as described for **A**). Means and standard deviations of at least three biological replicates are shown. Statistical significances were determined for various strains compared to STM WT, and between SIN 119944 and SIN 335-5. Student’s *t*-test was applied and results are indicated as: n.s., not significant; *, *p* < 0.05; **, *p* < 0.01; ***, *p* < 0.001.

To further test a correlation between presence of pESI and the atypical biofilm formation at high temperatures, further strains with and without pESI were investigated (**Figure S 1**). The megaplasmid pESI is transmitted by conjugation, and its acquisition by *S. enterica* serovar Muenchen (SMU) and other serovars has been reported (Cohen et al., 2022a; Dos Santos et al., 2022). While older SMU isolates in *Salmonella* reference collection B (SARB) lacking pESI did not produce atypical biofilm, a set of recent isolates harboring pESI all showed high levels of biofilms after culture at 42 °C for 24 h. Experimental conjugational transfer of pESI to recipient strains with suitable selectable markers was performed. Transfer of pESI did not confer atypical biofilm formation to *E. coli* ORN172. However, SIN 335-3 and STM harboring pESI produced atypical biofilm, although at levels lower than those of SIN 119944. We conclude that presence of pESI in SIN and other *S. enterica* serovars confers an atypical biofilm phenotype at conditions similar to the mammalian and avian body temperature, i.e. 37 °C and 42 °C, respectively.

*pESI-mediated atypical biofilm formation is independent of biofilm master regulator CsgD* We next investigated whether the master regulator CsgD controls atypical biofilm formation of SIN [pESI]. **Error! Reference source not found.** shows that biofilm formation of SIN Δ*csgD* at 30 °C was significantly decreased, but still significantly higher than STM Δ*csgD*. Moreover, while no biofilm formation was determined for STM WT and STM Δ*csgD* at 37 °C or 42 °C biofilm formation of SIN 119944 at 37 °C or 42 °C was not affected by deletion of *csgD*.

In addition to CsgD, various global regulatory mechanisms have been reported to affect biofilm formation in *E. coli* and *S. enterica* (Azriel et al., 2016; Gerstel & Römling, 2003; Latasa et al., 2012). Using a set of mutant strains of SIN 119944 defective in global regulators, we investigated the potential contribution to atypical biofilm formation (**Figure S 2**). We observed that the absence of *soxR*, *fliZ*, *lrp*, *ompR*, *oxyR*, *phoP*, *fur*, or *fnr* did not alter biofilm formation of SIN 119944 at 37 °C and 42 °C. Moderate reductions of biofilms were determined for SIN 119944 deficient in *arcA* or *arcB*, while highly reduced biofilm was formed by Δ*rpoS* or Δ*rpoE* strains. However, lack of RpoS or RpoE led to highly reduced growth in CFA broth, indicating that the effects on biofilm formation are indirect and caused by altered physiology of the mutant strains.

We conclude that in addition to the known biofilm formation controlled by CsgD, atypical biofilm formation at higher temperatures is mediated by factors encoded by pESI.

### The presence of pESI alters motility

Biofilm formation requires change from planktonic to sessile lifestyle of bacteria, and involves increased expression of adhesion factors, biofilm matrix components, and repression of motility. The adhesion to abiotic surfaces and motility as function of pESI were analysed on single cell level by live-cell microscopy (**Figure 2**). We observed per field of view appr. 200 non-motile bacteria of strain SIN 119944 adherent to glass surfaces, while only appr. 10 bacteria of strain SIN 335-3 were sessile under these conditions (**Figure 2**A). Quantification of the motile population (**Figure 2**B) revealed an inverse distribution, with about 90% of STM and SIN 335-3 in the motile fraction, whereas only ca. 5% of SIN 119944 and SIN Δ*flhDC* bacteria exhibited motility. The average velocity was highly reduced for SIN 119944 and SIN Δ*flhDC*, but the small fraction of motile SIN 119944 exhibited velocities in the same range as SIN 335-5 (**Figure 2**C). We concluded from these experiments that the presence of pESI may increase the frequency of non-motile bacteria in the SIN population.

**Figure 2.**
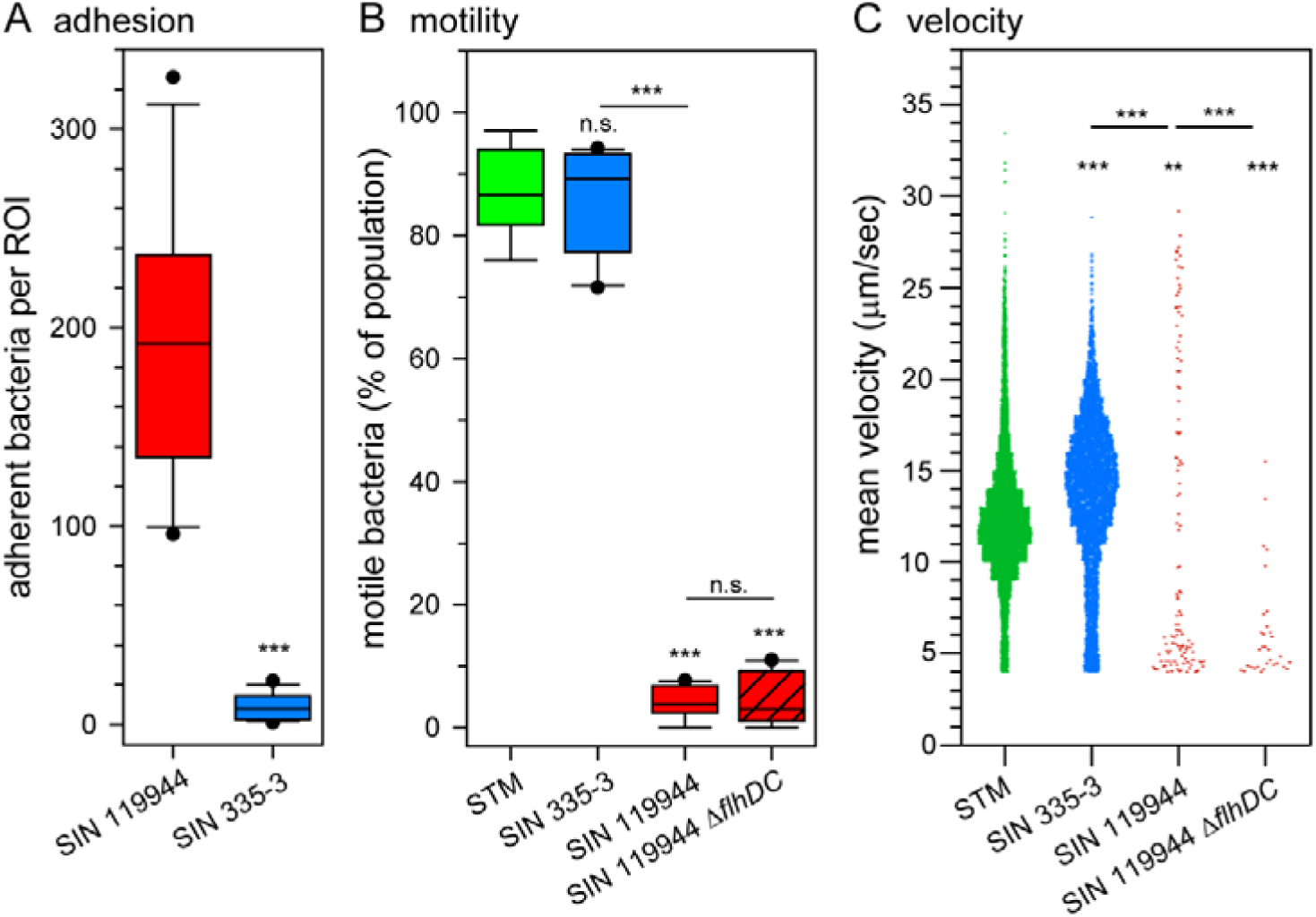
Emergent strain SIN 119944 exhibits increased adhesion to abiotic surfaces and reduced motility. STM WT, SIN 335-3, SIN 119944 and isogenic mutant strain Δ*flhDC* were analysed for adhesion to glass surfaces (**A**), motility (**B**), and velocity (**C**) of single bacterial cells. For motility, 12,094, 11,639, 3,203 and 2,375 individual bacteria of STM, SIN 335-3, SIN 119944, and SIN 119944 Δ*flhDC*, respectively, were analysed and results are indicated as: n.s., not significant; *, *p* < 0.05; **, *p* < 0.01; ***, *p* < 0.001.

### A novel autotransporter adhesin is present in biofilms formed by SIN 119944

We next set out to identify pESI-encoded factors contributing to the atypical biofilm formation. For this, unbiased bioinformatic mining and shotgun proteomics were performed. To identify candidate proteins with adhesive properties, the SPAAN prediction tool was applied to the predicted SIN 119944 proteome. This revealed a set of candidate proteins encoded by pESI, including the Klf and Ipf fimbriae, and the novel candidate GVI52_RS2310. Ipf and Klf are pESI-encoded fimbrial adhesins of the C/U pathway, and were already characterized (Aviv et al., 2017). SIN [pESI] mutant strains deficient in *klf*, *ipf*, or both *klf* and *ipf* were not affected in terms of the atypical biofilm phenotype (**Figure S 1**). GVI52_RS2310 yielded a SPAAN score (P_ad_) of 0.928, indicating a very high probability of adhesin function, for presence in the outer membrane, and is predicted as a putative β-barrel protein of 183 kDa anchored in the outer membrane.

We anticipated that proteins involved in atypical biofilm formation of SIN [pESI] would be more abundant in the proteome of SIN [pESI] forming biofilm compared to proteome of planktonic SIN [pESI]. Label-free mass spectrometry of bacteria in biofilm material versus planktonic bacteria indicated a set of candidate proteins that were more abundant in sessile SIN [pESI] (**Figure 3**A). Among pESI-encoded proteins, the most abundant in biofilm-forming bacteria was the GVI52_RS2310-encoded protein (**Figure 3**B). The ORF GVI52_RS2310 is located between the ORFs encoding a RfaH homolog and an EAL-domain-containing protein (**Figure 3**C).

**Figure 3.**
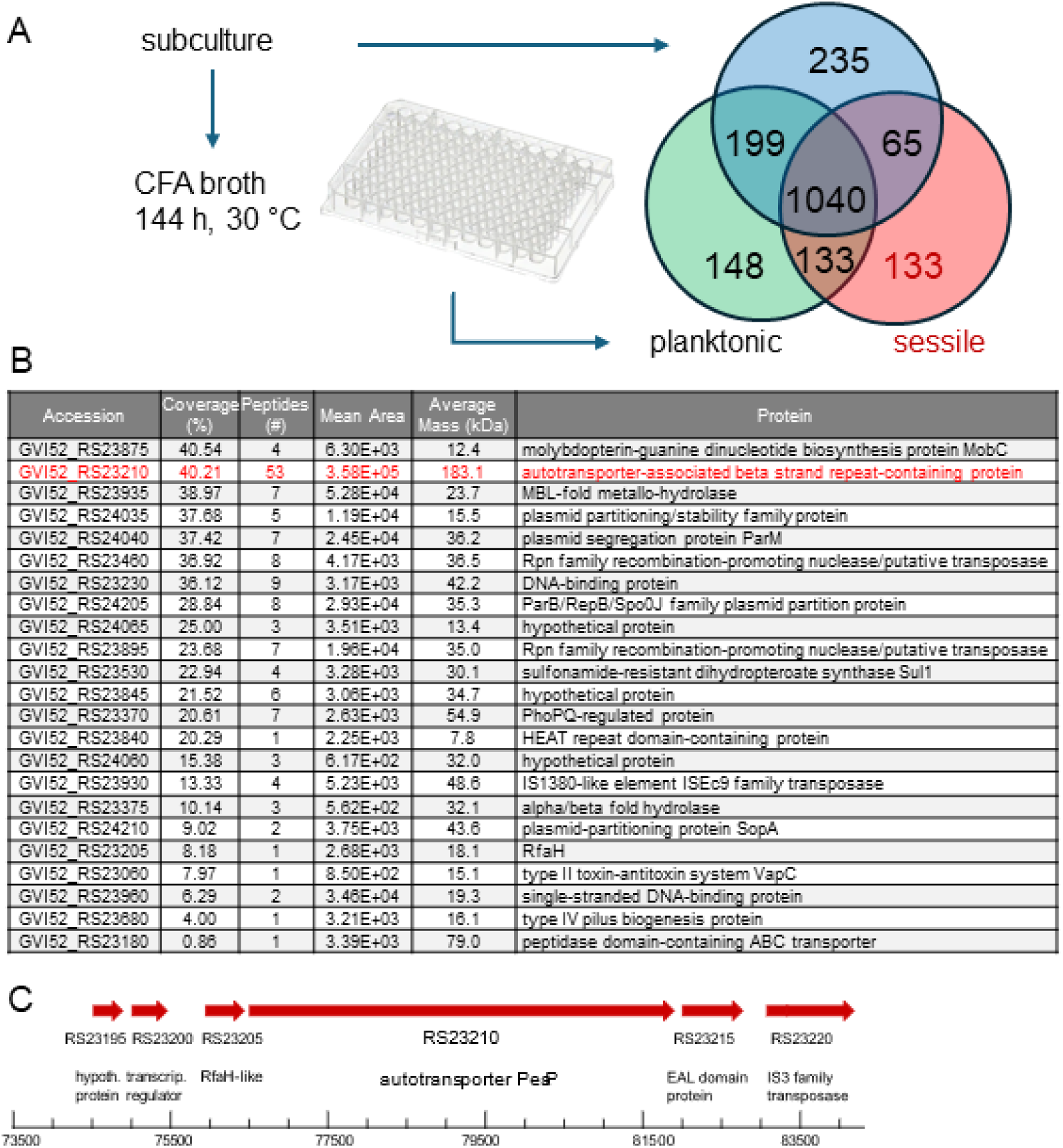
Proteomic analyses of biofilm components of SIN 119944 identified novel autotransporter PeaP. **A**) SIN 119944 was subcultured in CFA broth and further incubated in CFA broth at 30 °C in wells of microtiter plates. After incubation for 144 h, the supernatant was collected as fraction ‘**planktonic’**. After washing, adherent bacteria were recovered by scraping and analysed as fraction ‘**sessile’**. A portion of subculture was also analyzed as fraction ‘**subculture’**. Proteomes of the three fractions were analysed and numbers of proteins identified commonly or uniquely in the various fractions are indicated in the VENN diagram. Of the 133 proteins identified specifically in the proteome of the sessile population, proteins encoded by genes on pESI were selected in **B**). The most abundant pESI-encoded protein was GVI52_RS23210, a putative autotransporter of 183.1 kDa. We assigned the acronym PeaP to this protein. **C**) Genomic context of *peaP* on pESI. The ORF consists of 5,382 bp.

### PeaP is a novel autotransporter adhesin

Structural modeling of GVI52_RS2310 revealed the typical domain structure of an autotransporter protein (**Figure 4**). Accordingly, we suggest to term the GVI52_RS2310-encoded protein as ‘pESI-encoded autotransporter protein’ with the acronym PeaP. Domain organization of PeaP features a signal peptide (aa 1-23), a large passenger domain (aa 24-1478) and a β-barrel, typical for monomeric autotransporter adhesins (aa 1523-1793). The passenger domain and β-barrel are connected by a short α-helical portion (aa 1479-1522) predicted to be located within the β-barrel. The passenger domain comprises an N-terminal head domain with a disordered region (aa 24-446) and a C-terminal disordered region (aa 1383-1478). The central part of the passenger domain (aa 447-1382) shows a high β-sheet content. A remarkable feature of the passenger domain is the presence of three almost identical 101 aa repeats. The sub-domains aa 731-831 and aa 832-932 share an identical sequence, whereas the repeat aa 630-730 contains a single altered residue. This region is followed by an incomplete repeat of aa 731-769 that shares 97% identity. Interestingly, DNA sequences encoding the repeats show the same high degree of identity. This may indicate recent internal gene duplication [PMID: 15123416] events that were required to obtain sufficient size for the adhesin. We did not observe any potential sites for autoproteolytic processing within PeaP.

**Figure 4.**
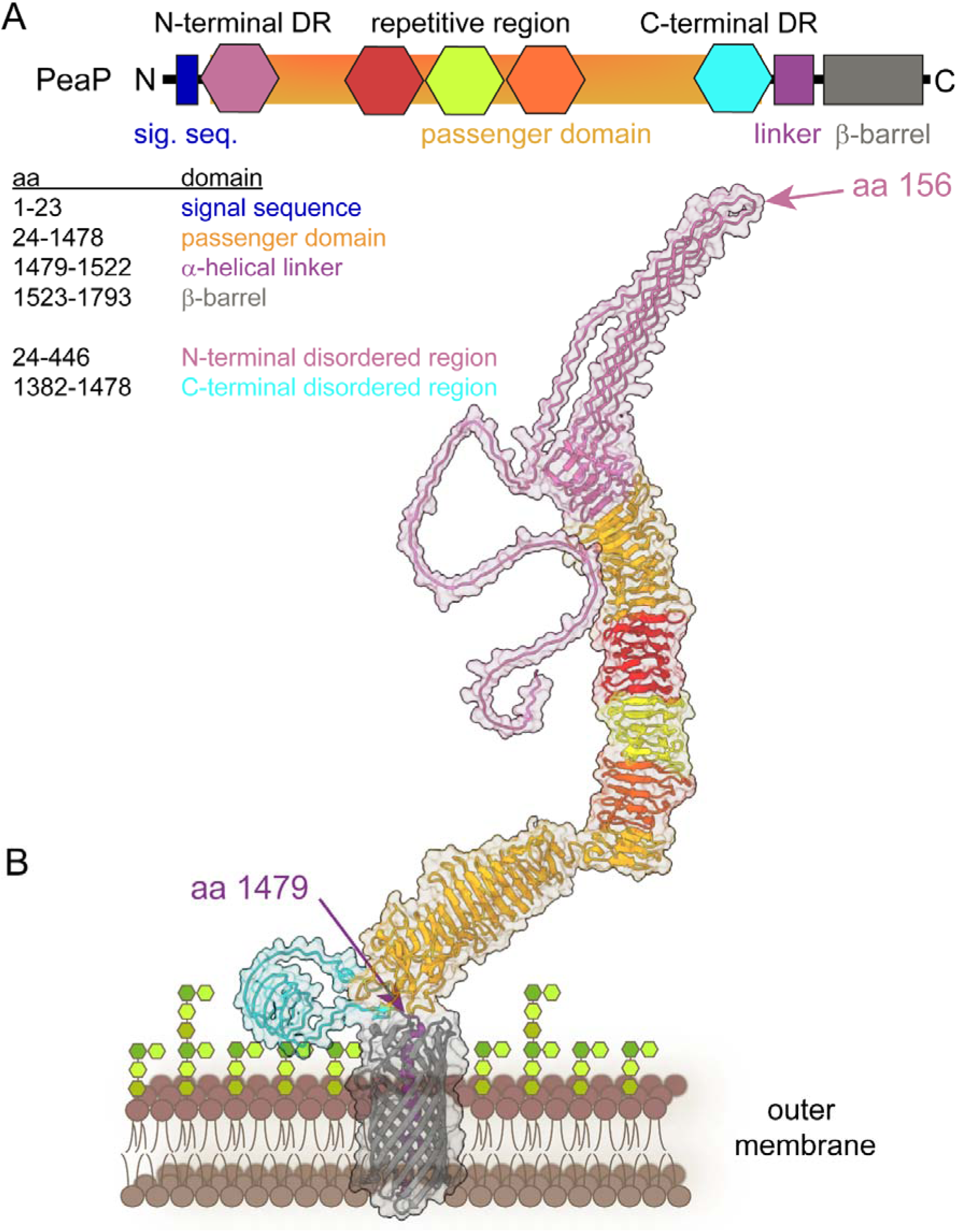
Domain organization and structure prediction of novel autotransporter PeaP. **A**) PeaP is predicted Sec system-secreted, and consists of a passenger domain (aa 22-1,478, yellow) and an outer membrane-integral β-barrel (aa 1,523-1,793, grey), connected by a short α-helical linker (aa 1,479-1,522, violet). **B**) The structure of PeaP was predicted using AlphaFold 3. The passenger domain features a N-terminal disordered region (aa 24-446, magenta) and a C-terminal disordered region (aa 1,382-1,478, teal). Analyses of sequence repeats revealed three repeat regions in the passenger domain (red, green, orange).

Database searches indicate that PeaP is only present in SIN and other *S. enterica* serovars harboring pESI. Thus, at present we cannot draw conclusions about the evolutionary origin of *peaP* or its integration into pESI. ProtBLAST/PSI-BLAST searches (Zimmermann et al., 2018) using the PeaP passenger domain indicated a number of bacterial outer transporters with similar domain organization, such as UniProt A0A839VPU9 of *Herbaspirillum* sp. Sphag64, UniProt A0A7K0Q0T5 of *Actinobacteria* bacterium, or UniProt A0A2N7X684 of *Trinickia symbiotica*. With 1,101, 1,195, and 1,236 aa, these proteins are significantly smaller than PeaP. *Herbaspirillum* sp. Sphag64 and *Trinickia symbiotica* are root-colonizing β-proteobacteria of the *Burkholderia* family, while *Actinobacteria* form a separate class of typically soil-inhabiting bacteria. The role of these autotransporter proteins in the biology of the indicated species is not known. In summary, PeaP is a new autotransporter adhesin acquired by *S. enterica* serovars via pESI.

### PeaP forms filamentous outer membrane appendages

PeaP is predicted as a monomeric outer membrane protein of 183 kDa, and we hypothesized that the mass and extended structure of the passenger domain may enable direct ultrastructural analyses. The immunolabeling shown in **Figure 6** indicated homogeneous distribution of PeaP on the surface of SIN, and accessibility for antibody binding. Thus, we set out to perform negative-stain transmission electron microscopy (TEM) to visualize PeaP (**Figure 5**). TEM of SIN 119944 revealed presence of numerous short filamentous surface structures that project from the outer membrane (**Figure 5**A). These structures were absent in SIN 119944 Δ*peaP,* SIN 335-3 and SMU lacking pESI (**Figure 5**BCF). In other *S. enterica* serovars harboring pESI such as SIN 335-3 [pESI] and SMU [pESI] (**Figure 5**DE), identical surface structures were identified.

**Figure 5.**
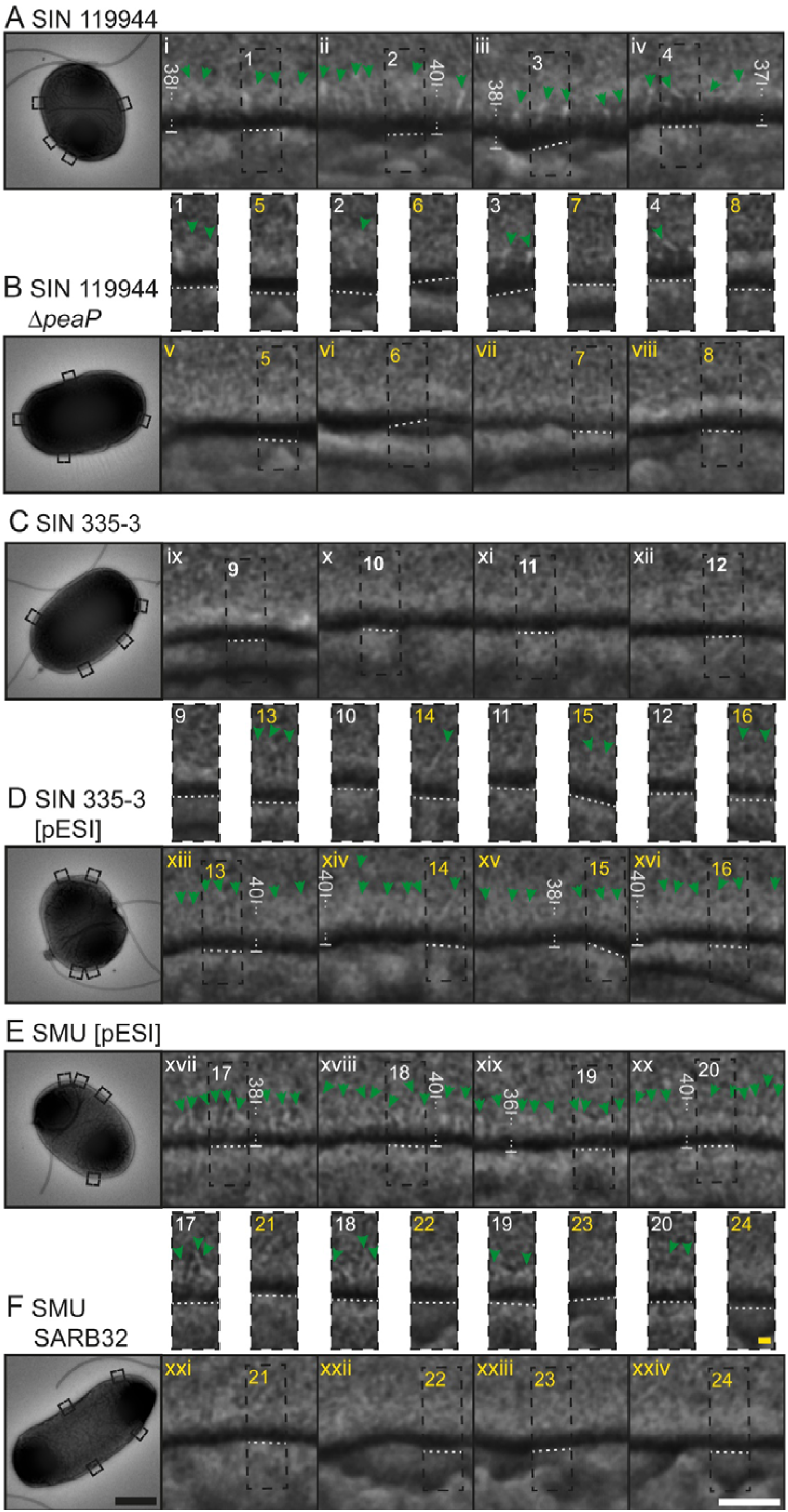
Autotransporter PeaP forms short filamentous appendages on the bacterial surface. PeaP surface expression was visualized by TEM of negative-stained samples. Strains *S*. Infantis 119944 (SIN, **A**), SIN 119944 Δ*peaP* (**B**), SIN 335-3 (**C**), SIN 335-3 [pESI] (**D**), *S*. Muenchen [pESI] (SMU, **E**), and SMU SARB32 (**F**) were cultured in CFA broth at 42 °C o/n with aeration and subcultured for 3.5 h under the same conditions. Bacteria were harvested by centrifugation, fixed with 4% PFA, and washed with HEPES buffer. Bacterial suspensions were spotted onto EM grids and subjected to negative staining using 2% PTA for 1 min. After washing and drying, samples were examined using TEM (Zeiss 912 TEM) operated at 80 keV. PeaP molecules appear as short (34-42 nm) filamentous protrusions in strains harboring pESI and intact *peaP* (details indexed white) and are indicated by green arrowheads. These structures are absent in the Δ*peaP* strain, pre-epidemic SIN 335-3, and SMU SARB32 (details indexed yellow). Scale bars: overview images (black bar), 500 nm; enlarged images (white bar), 50 nm; detail images (yellow bar), 10 nm. Measured distances from outer membrane periplasmic face and most distal part of PeaP are indicated in nm.

**Figure 6.**
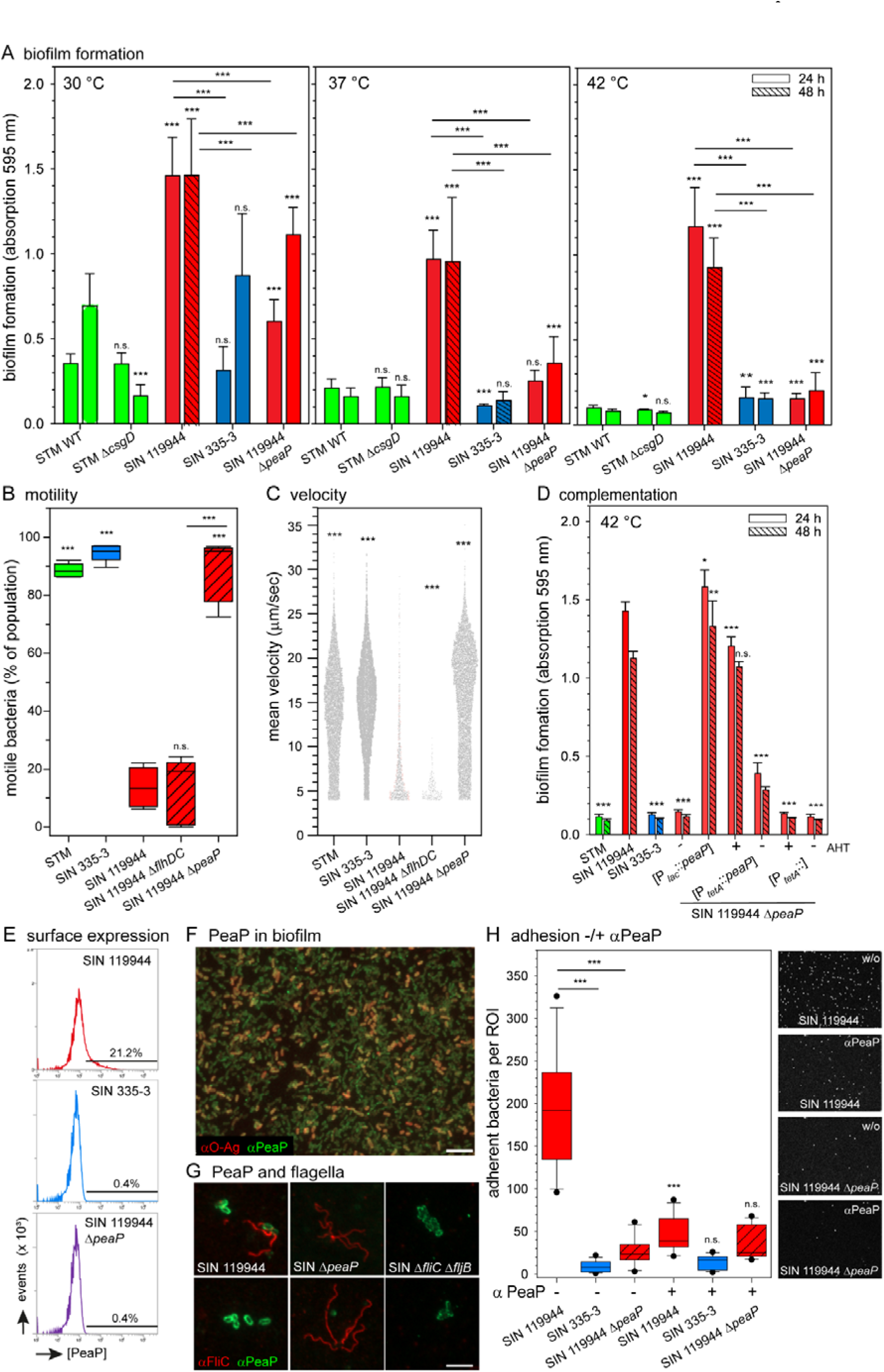
PeaP is required for biofilm formation, adhesion to abiotic surfaces, and sessile lifestyle of SIN. Biofilm formation on plastic surfaces (**A**), and motility (**B**) and velocity (**C**) on single cell level was compared for SIN 119944 WT to SIN Δ*peaP* and various strains as indicated. **D**) Atypical biofilm formation of SIN Δ*peaP* was complemented by plasmid-borne *peaP*. **E**) Antiserum raised against PeaP specifically detected surface-expressed PeaP on SIN 119944 after culture in CFA broth, and in biofilms formed on abiotic surfaces. **F**) Sessile SIN 119944 were immuno-stained for LPS O-Ag (red) and PeaP (green). **G**) SIN strains as indicated after culture in CFA broth were subjected to immuno-staining of PeaP (green) and flagellin FliC (red). **H**) Neutralization of PeaP-mediated adhesion by antibody. Various strains as indicated were subcultured, incubated in buffer without (-) or with (+) antiserum against PeaP, and added to wells of multi-chamber slides. After washing, numbers of adherent bacteria per fields of view were quantified. Statistical analyses based on at least three biological replicates were performed by ANOVA and results are indicated as: n.s., not significant; *, *p* < 0.05; **, *p* < 0.01; ***, *p* < 0.001.

The filamentous protrusions of PeaP showed an average extension of 37 nm, with a rather variable length distribution between 33 and 42 nm. PeaP molecules were found that project linear, as well as curved or bend. For some of the PeaP molecules the most distal portion appeared with wider diameter than the portion connecting to the outer membrane. However, the limited resolution of negative-stain TEM prevents further quantification of diameters. Some of the PeaP molecules appeared aggregated, especially in areas with a higher density of PeaP. The length of PeaP observed in ultrastructure by negative-stain TEM is in line with the predicted structure shown in **Figure 4**B, and the observed structural heterogeneity may a result of two extended disordered domains in PeaP (aa 24-446 and aa 1382-1478) leading to flexibility.

To further corroborate ultrastructural properties of PeaP, internal deletions were generated that removed predicted domains of PeaP, or parts of these (**Figure S 3**). The WT and mutant alleles of *peaP* were episomal expressed under control of the *tetR* P*_tetA_*cassette, and surface expression of PeaP was confirmed by flow cytometry (**Figure S 3**B). We also performed structure predictions for the deletion alleles of PeaP, generated models of outer membrane-anchored PeaP, and determined the distances from the outer membrane to the most distal portion of PeaP (**Figure S 3**C). Negative-stain TEM showed altered surface structures for the mutant alleles of PeaP (**Figure S 4**C). Analyses of PeaP length revealed 37.4 ± 2.4 nm for WT PeaP of SIN 119944 (means and standard deviations of at least 100 appendages), 37.2 ± 2.4 nm for WT PeaP expressed under control of *tetR* P*_tetA_*, 30.6 ± 2.2 nm for PeaP Δaa 27-446, 21.1 ± 1.4 nm (Δaa 447-1382), 32.0 ± 1.7 nm (Δaa 632-955), and 31.2 ± 2.3 nm (Δaa 1382-1478). For deletions within the passenger domain, the reduced size correlated with the extent of the deletion, and is in line with the predicted molecule structure.

### PeaP mediates atypical biofilm formation

To test the contribution of PeaP to the atypical biofilm formation by SIN, a mutant strain deleted for *peaP* in the background of SIN 119944 was generated and analysed. The Δ*peaP* strain was highly reduced in rapid biofilm formation at 42 °C, and biofilm amounts similar to STM or SIN 335-3 were determined (**Figure 6**A). The motility of PeaP-deficient SIN 119944 was similar to motility of STM WT and SIN 335-3, and velocities of motile cells were also found in the same range as observed for STM WT and SIN 335-3 (**Figure 6**BC). The specificity of the *peaP* deletion was confirmed by complementation with a plasmid harboring a WT allele of *peaP* that restored atypical biofilm formation (**Figure 6**D).

Native PeaP was recovered from protein extracts of SIN 119944 cultures and antiserum was raised in rabbits. Antibodies against PeaP specifically labelled SIN 119944 bacteria but not SIN 335-3 or SIN 119944 Δ*peaP* (**Figure 6**E). We used the antiserum to detect PeaP in biofilms formed by SIN 119944 at 42 °C and observed that the majority of sessile bacteria were positive for PeaP surface expression (**Figure 6**F). Co-labelling with antisera against PeaP and the O-antigen of LPS (O-Ag) indicated that surface expression of PeaP severely reduced labeling of the same cell by the anti-O-Ag antibody (**Figure 6**F). This partial exclusion suggests the PeaP covers the bacterial surface in a way that prevents antibody binding to O-Ag.

The highly reduced motility of SIN 119944 can either be explained by reduced expression of flagella, lack of flagella rotation, or strong adhesion that counteracts the force generated by rotating flagella. To test these possibilities, co-immunostaining for PeaP and flagella was performed (**Figure 6**G). Individual cells of SIN 119944 were positive for PeaP surface expression and also flagella. If bacteria were located in clusters, these were frequently devoid of immunolabeling of flagella. We observed that bacteria labelled by PeaP antibodies showed a homogeneous staining pattern that outlined the cell body. This is in contrast to the clustered distribution we previous observed for SiiE (Wagner et al., 2011), and again is indicative for a high density of PeaP molecules in the outer membrane.

To further test the contribution of PeaP to adhesion to abiotic surfaces, we neutralized PeaP by incubating *Salmonella* with αPeaP antiserum (**Figure 6**H). We determined that preincubation of SIN 119944 with αPeaP antiserum reduced the number of bacteria adhering to glass surfaces by a factor of 6.4, while no effect of the antiserum was observed for SIN 335-3 or SIN 119944 Δ*peaP*.

### PeaP increases the colonization of chicken gastrointestinal tract

To investigate potential contributions of PeaP to virulence, we investigated the invasion of the polarized epithelial cell line MDCK by SIN and STM (**Figure S 5**). Prior work revealed that adhesion factors such as SPI4-encoded SiiE are required to establish initial contact to the apical side of epithelial cells that then enables SPI1-T3SS-mediated invasion (Gerlach et al., 2008). Invasion of MDCK cells by SIN 119944 was 34.9-fold lower compared to STM WT, and invasion of pre-epidemic SIN 335-5 was even lower (593-fold lower than STM WT). The deletion of *peaP* in SIN 119944 had no significant effect on MDCK invasion. We therefore tested a potential contribution of PeaP in STM WT using the P*_tetA_*-controlled expression of *peaP*. The presence of plasmid p6443 for expression of *peaP* had no significant effect on MDCK invasion in the absence of the inducer AHT. When *peaP* expression was induced with AHT, invasion was reduced by a factor of 43. These data indicate that PeaP does not contribute to apical invasion of polarized epithelial cells, and that synthetic expression of *peaP* can interfere with STM adhesion and invasion mediated by expression of SPI1 and SPI4 genes and functions of their products. Initial tests with the chicken non-polarized epithelial cell line DF-1 did not indicate difference in invasion between SIN 119944 and SIN Δ*peaP*. The emergence of SIN [pESI] was associated with its presence in chicken flocks and with transmission to human hosts through poultry products (Alvarez et al., 2023; Gal-Mor et al., 2010). Given the physiological chicken body temperature of 40.6 to 41.7 °C and the dominance of PeaP-mediated adhesion and biofilm formation at 42 °C, we hypothesized that PeaP may contribute to colonization and/or persistence in chickens. Prior work demonstrated that colonization of the chicken intestinal tract by SIN [pESI] is increased compared to SIN lacking pESI (Drauch et al., 2021). Here, we focused on the contribution of PeaP and analysed SIN 119944 and its isogenic Δ*peaP* strain in a chicken infection model. The groups are shown in **Error! Reference source not found.**. Additionally, we included SIN strains constitutively expressing *lux* for bioluminescent detection of bacterial spread during infection. However, SIN strains lost the *lux* plasmid rapidly, and no antibiotic selection was possible for plasmid maintenance during infection. Thus, the results of groups 1 and 3 were combined as results for SIN 119944 WT infection and results of groups 2 and 4 were combined as SIN Δ*peaP*. Groups of 15 chickens were infected on day 2 after hatching and kept with additional 10 chickens which served as sentinels to observe horizontal transmission of SIN between chickens. Per group 3 infected and 2 sentinels were sacrificed on day 7, 14, 21, 28, and 35. Neither clinical symptoms and pathological results were found in any of the groups, nor differences in body weight, or weight of liver and spleen. The negative control group stayed negative throughout the animal trial. Cloacal swabs showed no significant difference in shedding between the groups, albeit 90% and 65% of the swabs taken from SIN 119944 WT or SIN Δ*peaP* infected birds at 1 d p.i., respectively, turned out positive. Thus, all animals, infected and sentinels, are combined per group and were considered infected. SIN was detected in the caecum of all infected birds, while only 26.26%, 39.39% and 32.32% of birds were positive in gallbladder, spleen or liver, respectively. SIN Δ*peaP* was lower in colonization of chicken gallbladder, spleen and liver compared to SIN 119944 (**Error! Reference source not found.**).

Quantitative analysis of colonization was conducted via CFU counts in caecum, gallbladder, spleen and liver (**Figure 7**). As our prior experiments indicated that surface expression of PeaP promotes adhesion to abiotic surfaces, as well as bacterial auto-aggregation, we decided to take these traits into account when estimating organ burden based on quantification of bacterial CFU. As PeaP is a proteinaceous factor, its proteolysis should alleviate adhesion and aggregation. Accordingly, we performed trypsin treatment to organ samples prior to plating for CFU determination. Comparison indicated significantly higher CFU counts after trypsin treatment in caecum and gallbladder (**Figure 7**), supporting that PeaP-mediated adhesion and potentially biofilm formation occurs in the chicken intestinal tract. Compared to SIN 119944, SIN Δ*peaP* showed significantly reduced colonization of caecum and gallbladder. We conclude that function of PeaP contributes to the efficient colonization of the chicken intestinal tract by epidemic pESI-positive SIN lineages.

**Figure 7.**
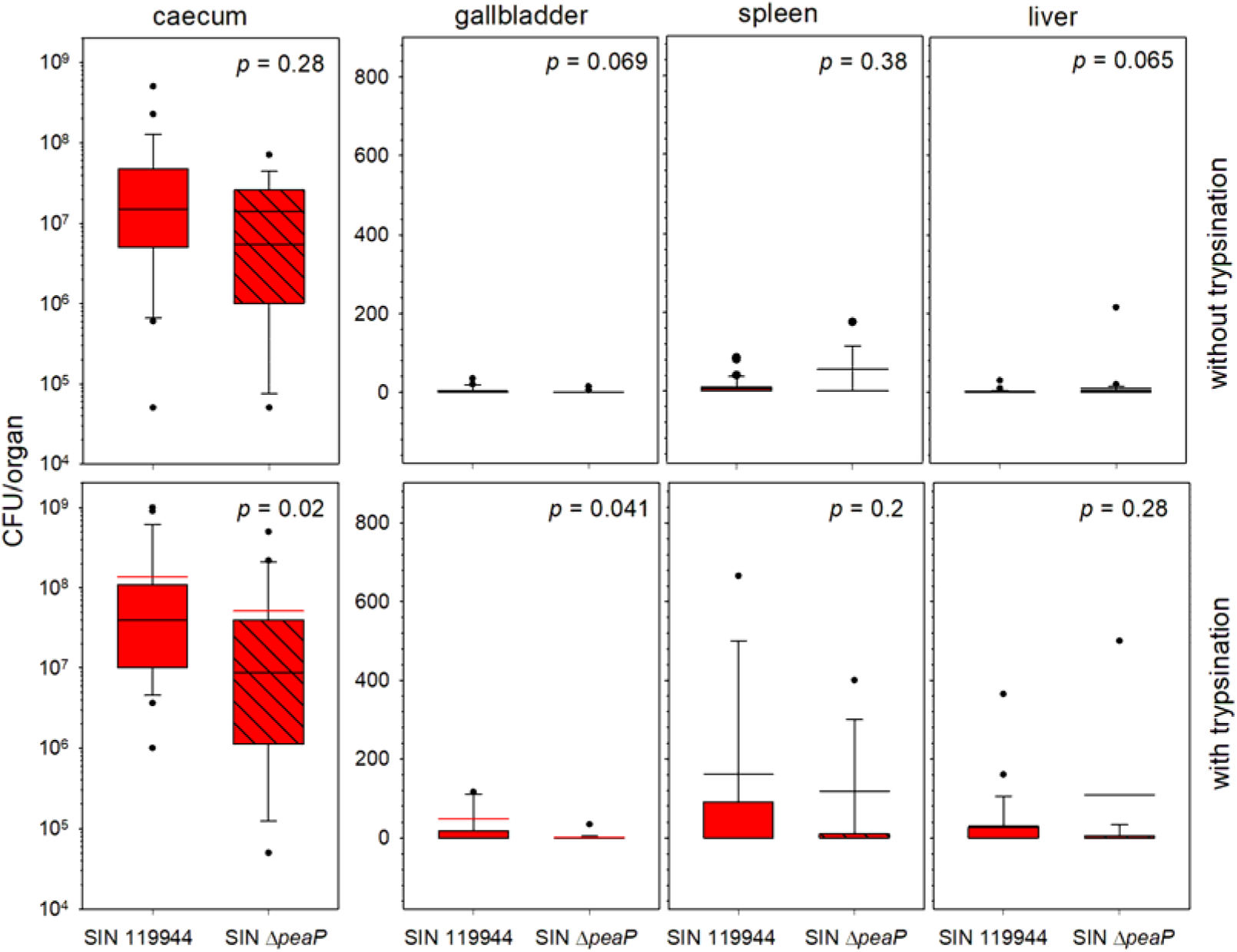
PeaP contributes to colonization of the chicken intestinal tract by SIN. 2-day-old Ross 308 broilers were infected with SIN 119944 or SIN Δ*peaP*. To determine the bacterial colonization in various organs, tissue homogenates of caecum, gallbladder, liver and spleen were plated on XLD agar and CFU counts were determined. Tissue homogenates were plated without trypsination, as well as with trypsination to release bacteria from aggregates prior to plating homogenates on agar plates for CFU determination. Trypsination was performed by incubation in 2.5 mg/ml Trypsin in PBS for 2 h at 37 °C with agitation at 300 rpm. Statistical analysis was performed using Wilcoxon test and significance values are indicated.

## Discussion

This work identified PeaP, a novel monomeric autotransporter adhesin mediating atypical biofilm formation in epidemic *S. enterica* serovar Infantis (SIN) isolates and other *S. enterica* strains harboring the megaplasmid pESI. PeaP-mediated adhesion and biofilm formation is independent from the central biofilm regulator CsgD and dominant at 42 °C, i.e. corresponding to avian body temperatures.

Strains of SIN harboring the megaplasmid pESI show remarkable success in global epidemic dissemination, especially in poultry (Alvarez et al., 2023; Mattock et al., 2024). Presence of pESI mediates multiple antibiotic resistances and encodes several fitness factors such as the complex biosynthesis, secretion, and uptake of the siderophore Yersiniabactin (Diamant et al., 2024). In addition to iron uptake, Yersiniabactin contributes to tolerance against oxidative stress in SIN.

Spread of pESI within *S. enterica* and to other enterobacteria occurs by conjugational transfer, and maintenance of the plasmid is ensured by multiple T/A systems encoded by pESI (Aviv et al., 2016; Cohen et al., 2022a). In addition to the large number of fimbrial adhesins encoded by operons on the chromosome and the virulence plasmid of various *S. enterica* serovars, we previously reported two pESI-encoded fimbrial adhesins, i.e. Ipf and Klp (Aviv et al., 2017). PeaP is the third pESI-encoded adhesin, and as an autotransporter adhesin adds to the family of autotransporters in *S. enterica* that includes MisL and ShdA as monomeric, and SadA as trimeric autotransporters. Using SPAAN bioinformatic analyses (Sachdeva et al., 2005), we recently identified further chromosomal-encoded autotransporter adhesin candidates in SIN such as BigA, YaiT and SinHI (Lüken et al., unpublished observation), but this approach did not indicate further pESI-encoded autotransporters other than PeaP.

The high surface expression level of PeaP under laboratory culture conditions is remarkable, and in contrast to most other members of the *S. enterica* adhesiome. For most adhesins, the environmental conditions leading to induction of gene expression and, consequently, detectable surface expression are still unknown. Accordingly, we previously devised a synthetic expression approach to control functional surface expression for analyses of adhesin structure and binding properties (Hansmeier et al., 2017). A reason for the tight control of expression of the canonical set of adhesins might be the integration in a complex regulatory network (reviewed in Blomfield & van der Woude, 2007). Adhesins are coregulated with expression of other biofilm matrix factors such Curli and cellulose synthesis (Gerstel & Römling, 2003), or coregulated with synergistically acting virulence factors such as SPI1-T3SS genes for host cell invasion and the *sii* operon encoding adhesin SiiE and (Gerlach et al., 2007; Main-Hester et al., 2008). This ensures that only one or few distinct adhesins are surface expressed under specific conditions. In contrast, *peaP* is a rather recent acquisition introduced by HGT of pESI, and it is likely that control of *peaP* expression is not fully integrated yet into the regulatory network responsible for the majority of adhesins in *S. enterica*. Klf and Ipf are further pESI-encoded fimbrial adhesins with distinct control of expression. For Klf, local control elements *klfB* and *klfL* and global control by Lrp were reported, while Ipf is under global control of Fur and OmpR (Aviv et al., 2017). We did not observed effects of deletions of *fur*, *ompR* or *lrp* on PeaP-dependent biofilm formation. This indicates that alternative, pESI-encoded regulators may be present that mediate control independent from the chromosomal regulators. Our bioinformatics analyses indicated at least 14 pESI-encoded proteins with potential function on control of gene expression. Two of these, a potential RfaH homolog (GVI52_RS23205) and an EAL domain-containing protein (GVI52_RS23215), are located upstream and downstream, respectively, of *peaP* and potentially co-transcribed. Further analyses will test a potential role in control of *peaP* expression. It will also be of interest to analyze if *peaP* expression interferes with other virulence functions such as SPI1/SPI4-mediated adhesion and invasion. Results of heterologous expression of *peaP* in STM (**Figure S 5**) point in such direction, but more detailed analyses in SIN are required to reveal potential interference.

Surface expression of adhesins is contributing to biofilm formation because they are components of the biofilm matrix. Important for conventional biofilm matrix formation are Curli fimbriae and Fim fimbriae. These factors are expressed under control of CsgD. The specific contribution of pESI to biofilm formation is masked by the conventional CsgD-controlled biofilm formation with cellulose and Curli as main matrix components. If CsgD-controlled biofilm formation is reduced such as during culture at 42 °C, PeaP-mediated biofilm formation is dominant. Here, we show that *peaP* expression is independent from CsgD function. So far, the mechanisms that regulate *peaP* are so far unknown, and central regulatory systems have minor effects on PeaP-mediated biofilm formation. PeaP appears as an autonomous factor for biofilm formation, because synthetic expression of *peaP* in STM leads to an atypical biofilm formation in heterologous strain backgrounds, although at lower level. The acquisition of pESI enabling expression three additional adhesins and biofilm formation at avian body temperature could explain the clonal success and rapid spread of SIN [pESI].

PeaP appears as novel bacterial adhesion factor, as no proteins with significant overall identity were found in database searches. Although PeaP has the typical architecture of an autotransporter adhesin with β-barrel, passenger, and head domains, the presence of two extended disordered regions in the N- and C-terminal portions of the passenger domain is remarkable. These regions may provide a high degree of flexibility for surface-expressed PeaP molecules as indicated by the heterogenous morphology of surface-expressed PeaP revealed by negative-stain TEM (**Figure 5**). Disordered regions in adhesins can also undergo conformational changes and adhesion-induced folding as exemplified for CdiA (Ruhe et al., 2018). Whether the disordered regions of PeaP share similar features, they need to be investigated by detailed structural analyses.

Of future interest will also be the characterization of binding properties and potential ligands for PeaP. The characterization of PeaP described here indicates a rather nonspecific, but tight interaction with abiotic surfaces such as glass cover slides and polystyrene of microtiter plates (**Figure 6**). PeaP-mediated adhesion is sufficiently strong to stop flagella-mediated motility as well as Brownian motion of bacterial cell bodies, and likely results from multiple adhesion events mediated by the large number of surface-expressed PeaP molecules, as observed in ultrastructural analyses. Whether additional specific ligands for PeaP exist is currently unknown, and will be investigated in our future work.

The epidemic spread of SIN harboring pESI in poultry, especially in broilers with subsequent transmission to humans, together with the observation of biofilm formation at 42 °C prompted us to investigate the importance of PeaP to colonize chicken. We determined that chicken infected with PeaP-positive SIN showed higher colonization of caecum and gallbladder, proving that PeaP supports colonization of host organisms and thus is expected to contribute to prolonged presence in animals. However, it is likely that additional pESI-encoded functions support efficient colonization and persistence, such as resistance against disinfectants, the Yersiniabactin iron uptake system with additional protection against oxidative stress (Diamant et al., 2024), and the multiple antimicrobial resistance factors. These factors are likely important in non-avian hosts, but in combination with PeaP as a potential host-colonization factor, they could provide SIN with the traits needed for epidemic success.

## Acknowledgements

This work was supported by grants of the German-Israel-Foundation to OGM and MH, and Hans Mühlenhoff-Stiftung for fellowship support of PF. MH acknowledges funding by the DFG in the framework of SFB 1557 P8 (467522186) and support by Z2 (467522186). We like to thank Monika Nietschke, Ursula Krehe, Delfina Jandreski-Cvetkovic and Claudia Ibesich for technical support, and Vesna Stanislajevic and Attila Sandor for maintenance of experimental animals. The support of Richard Liermann (chicken infection), Jörg Deiwick and Stefan Walter (MS analytics), and Katherina Psathaki (TEM imaging) is kindly acknowledged. We also acknowledge molecular graphics and analyses performed with UCSF ChimeraX, developed by the Resource for Biocomputing, Visualization, and Informatics at the University of California, San Francisco, with support from National Institutes of Health R01-GM129325 and the Office of Cyber Infrastructure and Computational Biology, National Institute of Allergy and Infectious Diseases.

## Materials and Methods

### Cultivation of bacterial strains

Bacterial strains used in this study are listed in **Table 1**. Routinely, bacteria were culture with aeration in LB or on LB agar at 37 °C if not otherwise stated. If necessary for selection of plasmids or resistances, carbenicillin (50 µg/ml), kanamycin (50 µg/ml), tetracycline (20 µg/ml), and/or chloramphenicol (12.5 µg/ml) were added to the media. For induction of the Tet-on system, overnight (o/n) cultures were diluted 1:31 in fresh medium in glass test tubes and grown aerobically by continuous rotation in a roller drum at 60 rpm for 1.5 h and anhydrotetracycline (AHT) was added to final concentration of 1 µg/ml and 10 ng/ml for SIN [pESI] and STM, respectively. Induced cultures were grown for additional 2 h before further analyses.

**Table 1.**
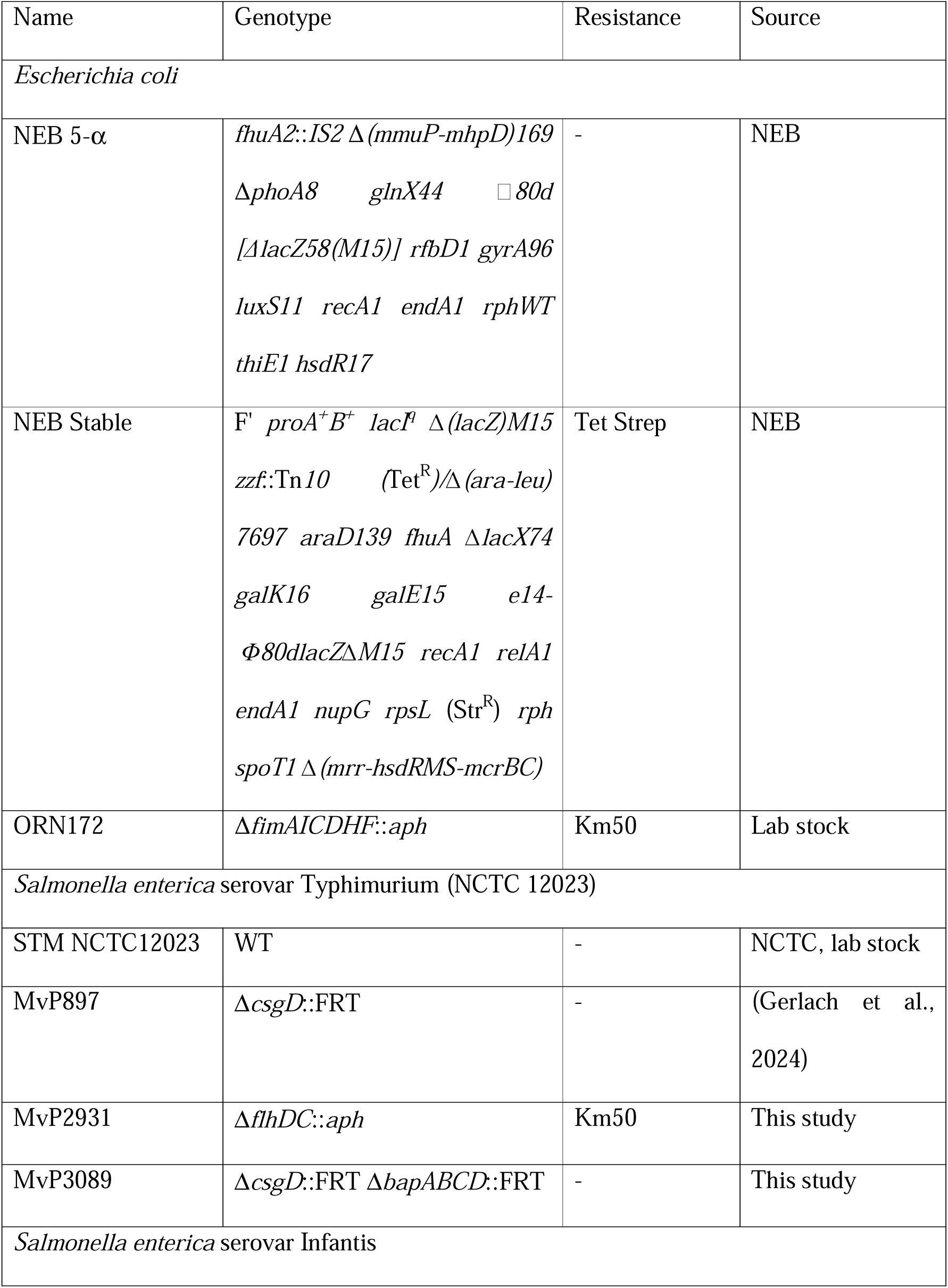
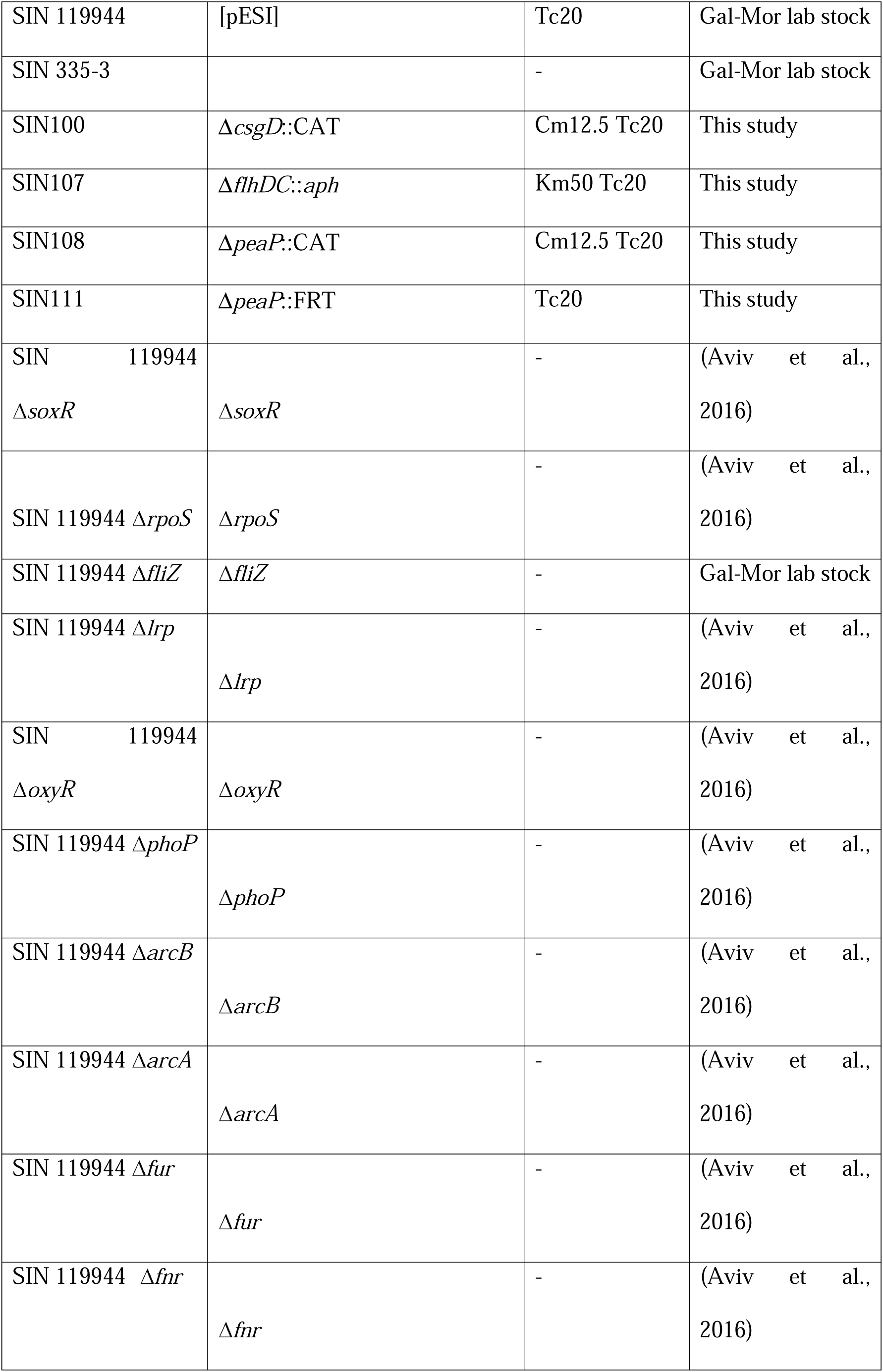
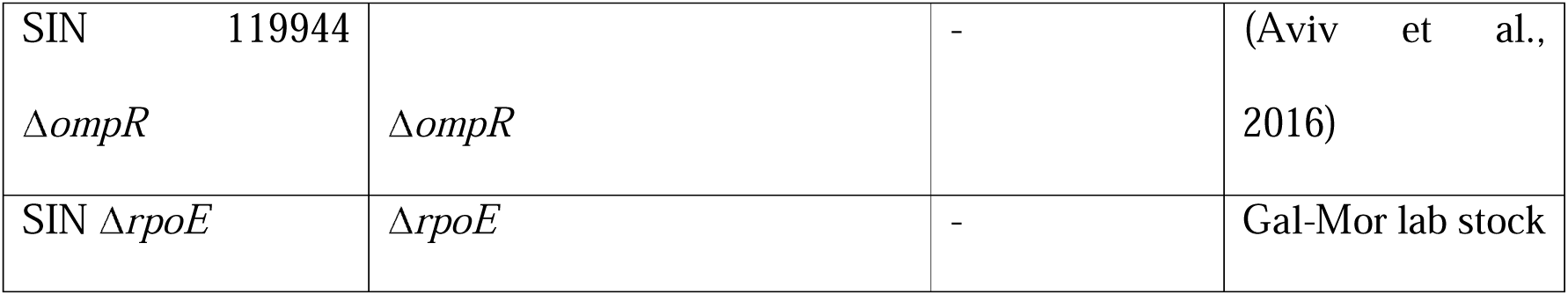
Bacterial strains used in this study.

### Generation of deletion alleles of peaP

Plasmids for expression of deletion alleles of *peaP* are listed in **Table 2** and were generated based on p6443. Deletions were generated by site-directed mutagenesis using the Q5 kit (NEB) according to manufacturers’ instructions with primers listed in **Table S 1.** Deletions were confirmed by control PCR and DNA sequencing, and synthesis of mutant alleles of PeaP by Western blot analyses with antiserum against PeaP.

**Table 2:** Plasmids used in this study.

| Plasmid | Relevant characteristics | Source |
| --- | --- | --- |
| pKD3 | template plasmid for CAT cassette, suicide vector (*ori* R6K), Cm<sup>r</sup> | (Datsenko & Wanner, 2000) |
| pKD46 | λ Red expression under control of arabinose-inducible promoter temperature sensitive, Carb<sup>r</sup> | (Datsenko & Wanner, 2000) |
| pE-FLP | FLP recombinase expression, Carb<sup>r</sup> | (St-Pierre et al., 2013) |
| pGEN-*lux* | *luxCDABE*, constitutive *lux* expression | Lab stock |
| p6443 | pWSK29 *tetR* P<sub>tetA</sub>::*peaP* | This study |
| p6581 | pWSK29 *peaP* | This study |
| p6660 | pWSK29 *lacI* P<sub>lac</sub>::*peaP* | This study |
| p6756 | pWSK29 *tetR* P<sub>tetA</sub>::*peaP* Δ aa27-446 | This study |
| p6757 | pWSK29 *tetR* P<sub>tetA</sub>::*peaP* Δ aa1382-1478 | This study |
| p6558 | pWSK29 *tetR* P<sub>tetA</sub>::*peaP* Δ aa447-1382 | This study |
| p6759 | pWSK29 *tetR* P<sub>tetA</sub>::*peaP* Δ aa632-955 | This study |

### Analysis of biofilm formation by crystal violet assay

For the quantification of biofilm formation, bacteria were cultivated o/n in colony-forming antigen (CFA) medium (1% Casamino acids, 0.15% Yeast extract, 50 mg/l MgSO_4_, 5 mg/l MnCl_2_, pH 7.4) and further subcultured in CFA medium as described before (Elpers & Hensel, 2020). Briefly, bacterial cultures were diluted to 3 x 10^8^ bacteria/ml, 150 µl bacterial suspension were filled per well in 96-well plates (Polystyrene, flat bottom without surface treatment, Hartenstein #MTD and #MTF) and plates were incubated in humid chambers for 24 h and 48 h at 30 °C, 37 °C and 42 °C. After incubation, OD_595_ was measured to control bacterial growth. Wells were washed three times with H_2_O_dd_ to remove planktonic bacteria. Biofilm material was stained using 0.1% crystal violet in H_2_O_dd_ for 5 min at RT. Wells were washed again three times with H_2_O_dd_, and the bound crystal violet stain was extracted by adding 150 µl EtOH abs. for 10 min at RT. Finally, the OD_595_ was measured.

### Analysis of motility of single cells

For the analysis of bacterial adhesion to plastic surfaces, bacteria were grown aerobically in LB medium o/n at 37 °C and further subcultured 1:31 in fresh LB medium for 3.5 h at 37 °C aerobically. If required, the Tet-On system was induced as described in ‘Cultivation of bacterial strainś. After subculture, OD_595_ was measured and bacteria diluted in PBS to an OD_595_=0.1 in PBS. 150 µl of bacterial solution was added to each well of an µ-Slide 8 Well (Ibidi #80806) and directly used for microscopy (Zeiss AxioObserver with Photometrics CoolSnap HQ^2^, 150 ms exposure, bright field illumination, 40x objective, 1 min video sequence). Video sequences were further analysed using ImageJ (FIJI, version 1.52c) and the plug-In “Track-Mate” with following settings: estimated blob diameter 5 µm, threshold 1.25-2.0, max. linking distance 10, Gap-closing distance 5, max. Gap-closing frame gap 3, ≥ 10 spots per track, ≥ 4 µm/sec mean velocity = swimming). At least three biological replicates with 3 video sequences were analysed.

### Analysis of adhesion to abiotic surfaces

For the analysis of bacterial adhesion to plastic surfaces, bacteria were grown aerobically in LB medium o/n at 37 °C and further subcultured 1:31 in fresh LB medium for 3.5 h at 37 °C aerobically. Induction of the Tet-On system took place as described in ‘Cultivation of bacterial strainś, if needed. After subculture, OD_595_ was measured and bacteria diluted in PBS to an OD_595_=0.1 in PBS. 150 µl of bacterial solution was added to each well of an µ-Slide 8 well (Ibidi #80806, Gräfelfing, Germany) and incubated for 10 min at RT. After incubation, wells were washed six times with 150 µl PBS to remove non-bound bacteria, finally 150 µl PBS was added to each well and bacteria were imaged using (Zeiss AxioObserver with Photometrics CoolSnap HQ^2^) with 40x objective and 120 ms exposure time. At least 3 biological replicates with 3 images per strain were further analyzed using ImageJ (version 1.52c) and the PlugIn “TrackMate” for counting of present bacteria per image with the settings: 5 microns estimated blob diameter and a threshold of maximum 1.5.

### Bioinformatics and structure prediction by AlphaFold 3

The prediction tool SPAAN (Sachdeva et al., 2005) obtained from https://sourceforge.net/projects/adhesin/files/SPAAN/ and installed under Ubuntu 19.10 equipped with C compiler was used to identify genes on pESI encoding putative adhesins. Protein FASTA file of the coding sequences of pESI was obtained from NCBI (CP047882.1 (Cohen et al., 2020) and used as the input file. Due to the plasmid size the threshold of protein considered putative adhesins was set to *P*_ad_ > 0.7, as used for whole genomes. Signal peptide and cleavage site prediction of PeaP was carried out using SignalP-5.0 (Almagro Armenteros et al., 2019).

Structure predictions of PeaP were generated using the AlphaFold Server powered by AlphaFold 3 (Abramson et al., 2024). Predicted protein structures were visualized using ChimeraX v1.10 (Meng et al., 2023). Further structure predictions and homology analyses were conducted using the MPI Bioinformatics Toolkits HHpred and HMMER (Gabler et al., 2020; Zimmermann et al., 2018).

### Transmission electron microscopy

Cultures of strains harboring plasmids for expression of *peaP* were induced with 1 µg/ml AHT and concentrated by centrifugation at 4,000 x g for 5 min. After fixation with 4% PFA for 30 min, the bacterial pellets were washed with HEPES buffer (20 mM HEPES, 150 mM NaCl, pH 7.4) and finally dissolved in 1 ml of HEPES. As a first step, 10 µl of the bacterial suspension was dropped onto glow-discharged formvar/carbon-coated TEM grids. After 1 min, the suspension was blotted off with filter paper, and this step was repeated twice. Three rounds of washing were performed by applying 4 µl of distilled water to the grid and immediately blotted off liquid. The grids were stained with 4 µl of 2% phosphotungstic acid diluted in distilled water adjusted to pH 7.4 for 45 s, then the suspension was blotted off with a filter paper. Immediately after blotting, 4 µl of distilled water was applied, directly blotted off and the grids were air dried for 5 min. Transmission electron microscopy analysis was performed using a Zeiss Leo 912 system equipped with an in-column OMEGA energy filtering system and a Koehler high-contrast illumination system operating at 80□keV. Images were taken at 8,000x magnification and then processed in Photoshop: brightness, contrast and gamma were adjusted, a Gaussian-blur mask was applied, and a high-pass filter was used.

### Chicken infection model

A total of 115 one-day old ROSS 308 chicken (Brueterei Schulz, Lassnitzhoehe, Austria) were assigned to five groups: 25 birds per infected group and 15 birds in the negative control group (Table S 2). Each group was housed separately in an isolator (Montair HM2500, Montair Environmental Solutions B.V., Kronenberg, The Netherlands). Water and feed were provided *ad libitum*. Upon arrival, each bird was marked individually with a subcutaneous tag (Swiftag, Heartland Animal Health Inc. Fair Play, MO, USA).

For preparation of inoculum, SIN strains were cultured on MacConkey agar at 37 °C, aerobic for 24 h. A single colony was transferred into Luria-Bertani-Broth (LB, Invitrogen, Vienna, Austria) then incubated at 37 °C, aerobic, for 24 h in a shaking incubator (250 rpm). CFU counts were determined by plating on MacConkey agar in duplicate, allowing for the adjustment of the inoculum to a final concentration of 1 x 10□ CFU/ml.

The experimental design and sample collection was as follows: Birds were infected on day 2 of life via crop tube administration of 1 x 10□ CFU of the corresponding SIN inoculum. In each infected group, 15 birds were orally infected, while the remaining 10 birds served as sentinels and received 1 ml of phosphate-buffered saline (PBS, Gibco, Paisley, UK). All birds in the negative control group were also administered 1 ml of PBS (**Table S 2**).

Clinical parameters and housing conditions, including temperature, humidity, air pressure, and airflow, were monitored daily. Before infection and after infection cloacal swabs were collected on a weekly basis.

On days 7, 14, 21, 28, and 35, five birds (three infected and two sentinels) from each infected group and three birds from the negative control group were euthanized via intramuscular injection of a combination of Sedaxylan® (20 mg/ml, Dechra Pharmaceuticals, Dornbirn, Austria) and Narketan® (100 mg/ml, Vetoquinol, Vienna, Austria), followed by bleeding (*V. jugularis*). Necropsy was performed according to a standard protocol, recording body weight, liver and spleen weights, and gross pathological lesions. Tissue samples from the caecum, gallbladder, liver, and spleen were collected for bacteriological analysis.

Furthermore, paired cloacal swabs were analyzed to determine shedding patterns. One swab was directly streaked onto xylose-lysine-deoxycholate agar (XLD, Merck, Vienna, Austria) and incubated at 37 °C for 24 h. The second swab was stored at 4 °C and, in case of a negative direct plating result, subjected to an enrichment procedure following EN ISO 6579-1:2017 (EN ISO 6579-1:2017+A1:2020. Microbiology of the Food Chain – Horizontal Method for the Detection, Enumeration, and Serotyping of *Salmonella* – Part 1: Detection of *Salmonella* spp.).

To assess SIN colonization in organs, 1 g of liver, spleen, and caecum, as well as the entire gallbladder, were homogenized in PBS (Ultra Turrax T 10 basic, IKA, Staufen, Germany) and plated on XLD in 1:10 dilutions in duplicate. Trypsin (Merck, Darmstadt, Germany) was added to each sample to 2.5 mg/ml final concentration, followed by incubation at 37 °C for 2 h with agitation (300 rpm) before plating on XLD in 1:10 dilutions in duplicate (**Table S 3**).

Ethical approval for the animal infection trial was given by the institutional ethics committee and licensed by the national authority according to the Austrian law for animal experiments (license number GZ.: 2024-0.361.835).

Data were analysed with R (R Core Team 4.2.2) and exploratory data analysis was performed. Normality was assessed using the Anderson-Darling-Test. CFU data was not normally distributed, data was logarithmic transformed and Wilcoxon test was used to test for statistical significance between the two groups.

**Suppl. Material captions:**

**Table S 1. Oligonucleotides used in this study.**

**Table S 2. Design of the chicken infection experiment.**

**Table S 3. SIN detection in organs of infected chicken.**

**Figure S 1. Presence of pESI mediates atypical biofilm formation.** Various *S. enterica* isolates of serovars Typhimurium (STM), Infantis (SIN), Muenchen (SMU), and *E. coli* (EC) ORN172 and isogenic mutant strains were analysed for biofilm formation at 42 °C. The absence or presence of pESI is indicated by – and +, respectively. Biofilm formation was quantified as described for Figure **1**. Means and standard deviations of at least three biological replicates are shown. Statistical analyses for comparison to SIN 119944 were performed by ANOVA and results are indicated as: *, *p* < 0.05; **, *p* < 0.01; ***, *p* < 0.001, all other data are not significantly different (not indicated).

**Figure S 2. Role of global regulatory systems in atypical biofilm formation of SIN 119944.** Biofilm formation of STM WT, STM Δ*csgD*, SIN 119944, and various mutant strains isogenic to SIN 119944 as indicated was analysed as described for Figure **1**. Means and standard deviations of at least three biological replicates are shown. Statistical analyses for comparison to STM WT or SIN 119944 were performed by ANOVA and results are indicated as: *, *p* < 0.05; **, *p* < 0.01; ***, *p* < 0.001, all other data are not significantly different (not indicated).

**Figure S 3. Deletion of domains of PeaP and surface expression of PeaP alleles**. **A**) Domain organization of PeaP and position of domain deletions analysed in this study. Deletions in PeaP comprised aa 27-466 (N-terminal disordered region), aa 1,382-1,478 (C-terminal disordered region), aa 447-1,382 (passenger domain), aa 632-955 (passenger domain). **B**) Flow cytometry analyses of surface expression of WT PeaP and various mutant alleles. Surface expression was analyzed in SIN 119944 Δ*peaP* complemented with various plasmids as indicated. The tet-on expression system was used for episomal expression of WT *peaP* and mutant alleles of *peaP* with internal deletions as indicated. Strains were subcultured with (+) or without (-) addition of AHT, fixed, subjected to labelling and with antibodies against PeaP, and analysed by flow cytometry to quantify the percentage of PeaP-positive bacteria. **C**) The structures of PeaP and various deletion alleles were predicted using AlphaFold 3. The calculated distance between the α-helical linker (aa 1,479 in WT PeaP) and distal part of PeaP is indicated in nm.

**Figure S 4. Surface expression of truncated forms of PeaP**. SIN 119944 Δ*peaP* harboring the empty vector [vector] (**A**), or plasmids for AHT-inducible expression of WT *peaP* [*peaP*] (**B**) or deletion alleles of *peaP* as indicated (**C, D, E, F**) were used. Strains were cultured as described for **Figure 5**, and P*_tetA_*-controlled expression of *peaP* was induced by addition of 1 µg/ml AHT after subculture for 1 h. Negative-stain TEM of various strains was performed as for **Figure 5**. Green arrowheads indicate representative PeaP molecules. Scale bars: overview images (black bar), 500 nm; enlarged images (white bar), 50 nm; detail images (yellow bar), 10 nm.

**Figure S 5. PeaP is dispensable for invasion of polarized epithelial cells.** The polarized epithelial cell line MDCK was used as host cells for invasion by STM or SIN strains as indicated. STM harboring a plasmid for synthetic expression of *peaP* was used after subculture without or with addition of AHT for expression of *peaP* under control of P*_tetA_*. After infection for 60 min., cells were washed and medium containing 100 µg/ml Gentamicin was added to kill non-internalized bacteria. Infected MDCK cells were lysed and lysates plated for CFU determination. Invasion is expressed as percentage of internalized inoculum, and means and standard deviations of three replicates are shown.

**Movie S1. Presence of PeaP interferes with flagella-mediated motility of SIN.** Time lapse microscopy was performed for STM WT, and SIN 119944 WT, Δ*flhDC* or Δ*peaP* as indicated in the movie sections. Sequences of 60 sec were recorded by brightfield microcopy (Zeiss AxioObserver) using a 40x objective and DIC illumination. Scale bars, 20 µm.

## References

1. Abramson, J., Adler, J., Dunger, J., Evans, R., Green, T., Pritzel, A., Ronneberger, O., Willmore, L., Ballard, A. J., Bambrick, J., Bodenstein, S. W., Evans, D. A., Hung, C. C., O’Neill, M., Reiman, D., Tunyasuvunakool, K., Wu, Z., Zemgulyte, A., Arvaniti, E.,…Jumper, J. M. (2024). Accurate structure prediction of biomolecular interactions with AlphaFold 3. Nature, 630(8016), 493–500. 10.1038/s41586-024-07487-w

2. Alba, P., Leekitcharoenphon, P., Carfora, V., Amoruso, R., Cordaro, G., Di Matteo, P., Ianzano, A., Iurescia, M., Diaconu, E. L., Study Group, E. N., Pedersen, S. K., Guerra, B., Hendriksen, R. S., Franco, A., & Battisti, A. (2020). Molecular epidemiology of Salmonella Infantis in Europe: insights into the success of the bacterial host and its parasitic pESI-like megaplasmid. Microb Genom, 6(5). 10.1099/mgen.0.000365

3. Almagro Armenteros, J. J., Tsirigos, K. D., Sonderby, C. K., Petersen, T. N., Winther, O., Brunak, S., von Heijne, G., & Nielsen, H. (2019). SignalP 5.0 improves signal peptide predictions using deep neural networks. Nat Biotechnol, 37(4), 420–423. 10.1038/s41587-019-0036-z

4. Alvarez, D. M., Barron-Montenegro, R., Conejeros, J., Rivera, D., Undurraga, E. A., & Moreno-Switt, A. I. (2023). A review of the global emergence of multidrug-resistant Salmonella enterica subsp. enterica Serovar Infantis. Int J Food Microbiol, 403, 110297. 10.1016/j.ijfoodmicro.2023.110297

5. Aviv, G., Elpers, L., Mikhlin, S., Cohen, H., Vitman Zilber, S., Grassl, G. A., Rahav, G., Hensel, M., & Gal-Mor, O. (2017). The plasmid-encoded Ipf and Klf fimbriae display different expression and varying roles in the virulence of Salmonella enterica serovar Infantis in mouse vs. avian hosts. PLoS Pathog, 13(8), e1006559. 10.1371/journal.ppat.1006559

6. Aviv, G., Rahav, G., & Gal-Mor, O. (2016). Horizontal Transfer of the Salmonella enterica Serovar Infantis Resistance and Virulence Plasmid pESI to the Gut Microbiota of Warm-Blooded Hosts. mBio, 7(5). 10.1128/mBio.01395-16

7. Aviv, G., Tsyba, K., Steck, N., Salmon-Divon, M., Cornelius, A., Rahav, G., Grassl, G. A., & Gal-Mor, O. (2014). A unique megaplasmid contributes to stress tolerance and pathogenicity of an emergent Salmonella enterica serovar Infantis strain. Environ Microbiol, 16(4), 977–994. 10.1111/1462-2920.12351

8. Azriel, S., Goren, A., Rahav, G., & Gal-Mor, O. (2016). The Stringent Response Regulator DksA Is Required for Salmonella enterica Serovar Typhimurium Growth in Minimal Medium, Motility, Biofilm Formation, and Intestinal Colonization. Infect Immun, 84(1), 375–384. 10.1128/IAI.01135-15

9. Blomfield, I., & van der Woude, M. (2007). Regulation of Fimbrial Expression. EcoSal Plus, 2(2). 10.1128/ecosal.2.4.2.2

10. Brombacher, E., Baratto, A., Dorel, C., & Landini, P. (2006). Gene expression regulation by the Curli activator CsgD protein: modulation of cellulose biosynthesis and control of negative determinants for microbial adhesion. J. Bacteriol., 188(6), 2027–2037. http://www.ncbi.nlm.nih.gov/entrez/query.fcgi?cmd=Retrieve&db=PubMed&dopt=Citation&list_uids=16513732

11. Cohen, E., Kriger, O., Amit, S., Davidovich, M., Rahav, G., & Gal-Mor, O. (2022a). The emergence of a multidrug resistant Salmonella Muenchen in Israel is associated with horizontal acquisition of the epidemic pESI plasmid. Clin Microbiol Infect. 10.1016/j.cmi.2022.05.029

12. Cohen, E., Kriger, O., Amit, S., Davidovich, M., Rahav, G., & Gal-Mor, O. (2022b). The emergence of a multidrug resistant Salmonella Muenchen in Israel is associated with horizontal acquisition of the epidemic pESI plasmid. Clin Microbiol Infect, 28(11), 1499 e1497–1499 e1414. 10.1016/j.cmi.2022.05.029

13. Cohen, E., Rahav, G., & Gal-Mor, O. (2020). Genome Sequence of an Emerging Salmonella enterica Serovar Infantis and Genomic Comparison with Other S. Infantis Strains. Genome Biol Evol, 12(3), 151–159. 10.1093/gbe/evaa048

14. Datsenko, K. A., & Wanner, B. L. (2000). One-step inactivation of chromosomal genes in *Escherichia coli* K-12 using PCR products. Proc. Natl. Acad. Sci. U S A, 97(12), 6640–6645. 10829079

15. Diamant, I., Adani, B., Sylman, M., Rahav, G., & Gal-Mor, O. (2024). The transcriptional regulation of the horizontally acquired iron uptake system, yersiniabactin and its contribution to oxidative stress tolerance and pathogenicity of globally emerging salmonella strains. Gut microbes, 16(1), 2369339. 10.1080/19490976.2024.2369339

16. Dos Santos, A. M. P., Panzenhagen, P., Ferrari, R. G., & Conte-Junior, C. A. (2022). Large-scale genomic analysis reveals the pESI-like megaplasmid presence in Salmonella Agona, Muenchen, Schwarzengrund, and Senftenberg. Food microbiology, 108, 104112. 10.1016/j.fm.2022.104112

17. Drauch, V., Kornschober, C., Palmieri, N., Hess, M., & Hess, C. (2021). Infection dynamics of Salmonella Infantis strains displaying different genetic backgrounds - with or without pESI-like plasmid - vary considerably. Emerg Microbes Infect, 10(1), 1471–1480. 10.1080/22221751.2021.1951124

18. Elpers, L., & Hensel, M. (2020). Expression and functional characterization of various chaperon-usher fimbriae, Curli fimbriae, and Type 4 pili of enterohemorrhagic *Escherichia coli* O157:H7 Sakai. Frontiers in microbiology, 11, 378. 10.3389/fmicb.2020.00378

19. Espinoza-Erazo, V. P., Vela-Chauvin, M. G., Collantes-Vela, J. C., Zapata-Mena, S., & Machado, A. (2025). Biofilms of Salmonella: Implications for Food Safety and Public Health. Foodborne pathogens and disease. 10.1177/15353141251389597

20. Gabler, F., Nam, S. Z., Till, S., Mirdita, M., Steinegger, M., Soding, J., Lupas, A. N., & Alva, V. (2020). Protein Sequence Analysis Using the MPI Bioinformatics Toolkit. Curr Protoc Bioinformatics, 72(1), e108. 10.1002/cpbi.108

21. Gal-Mor, O., Valinsky, L., Weinberger, M., Guy, S., Jaffe, J., Schorr, Y. I., Raisfeld, A., Agmon, V., & Nissan, I. (2010). Multidrug-resistant Salmonella enterica serovar Infantis, Israel. Emerg Infect Dis, 16(11), 1754–1757. 10.3201/eid1611.100100

22. Gerlach, R. G., Claudio, N., Rohde, M., Jäckel, D., Wagner, C., & Hensel, M. (2008). Cooperation of *Salmonella* pathogenicity islands 1 and 4 is required to breach epithelial barriers. Cell. Microbiol., 10(11), 2364–2376. http://www.ncbi.nlm.nih.gov/entrez/query.fcgi?cmd=Retrieve&db=PubMed&dopt=Citation&list_uids=18671822

23. Gerlach, R. G., Jäckel, D., Geymeier, N., & Hensel, M. (2007). *Salmonella* pathogenicity island 4-mediated adhesion is coregulated with invasion genes in *Salmonella enterica*. Infect. Immun., 75(10), 4697–4709. http://www.ncbi.nlm.nih.gov/entrez/query.fcgi?cmd=Retrieve&db=PubMed&dopt=Citation&list_uids=17635868, http://pubmedcentralcanada.ca/picrender.cgi?artid=627567&blobtype=pdf

24. Gerlach, R. G., Wittmann, I., Heinrich, L., Pinkenburg, O., Meyer, T., Elpers, L., Schmidt, C., Hensel, M., & Schnare, M. (2024). Subversion of a family of antimicrobial proteins by *Salmonella enterica*. Front Cell Infect Microbiol, 14, 1375887. 10.3389/fcimb.2024.1375887

25. Gerstel, U., & Römling, U. (2003). The *csgD* promoter, a control unit for biofilm formation in *Salmonella typhimurium*. Res Microbiol, 154(10), 659–667. 14643403

26. Guzinski, J., Potter, J., Tang, Y., Davies, R., Teale, C., & Petrovska, L. (2023). Geographical and temporal distribution of multidrug-resistant Salmonella Infantis in Europe and the Americas. Frontiers in microbiology, 14, 1244533. 10.3389/fmicb.2023.1244533

27. Han, J., Aljahdali, N., Zhao, S., Tang, H., Harbottle, H., Hoffmann, M., Frye, J. G., & Foley, S. L. (2024). Infection biology of Salmonella enterica. EcoSal Plus, 12(1), eesp00012023. 10.1128/ecosalplus.esp-0001-2023

28. Hansmeier, N., Miskiewicz, K., Elpers, L., Liss, V., Hensel, M., & Sterzenbach, T. (2017). Functional expression of the entire adhesiome of *Salmonella enterica* serotype Typhimurium. Scientific reports, 7(1), 10326. 10.1038/s41598-017-10598-2

29. Latasa, C., Garcia, B., Echeverz, M., Toledo-Arana, A., Valle, J., Campoy, S., Garcia-Del Portillo, F., Solano, C., & Lasa, I. (2012). Salmonella Biofilm Development Depends on the Phosphorylation Status of RcsB. Journal of bacteriology, 194(14), 3708–3722. 10.1128/JB.00361-12

30. Liu, Z., Niu, H., Wu, S., & Huang, R. (2014). CsgD regulatory network in a bacterial trait-altering biofilm formation. Emerg Microbes Infect, 3(1), e1. 10.1038/emi.2014.1

31. Main-Hester, K. L., Colpitts, K. M., Thomas, G. A., Fang, F. C., & Libby, S. J. (2008). Coordinate regulation of *Salmonella* pathogenicity island 1 (SPI1) and SPI4 in *Salmonella enterica* serovar Typhimurium. Infect. Immun., 76(3), 1024–1035. http://www.ncbi.nlm.nih.gov/entrez/query.fcgi?cmd=Retrieve&db=PubMed&dopt=Citation&list_uids=18160484

32. Mattock, J., Chattaway, M. A., Hartman, H., Dallman, T. J., Smith, A. M., Keddy, K., Petrovska, L., Manners, E. J., Duze, S. T., Smouse, S., Tau, N., Timme, R., Baker, D. J., Mather, A. E., Wain, J., & Langridge, G. C. (2024). A One Health Perspective on Salmonella enterica Serovar Infantis, an Emerging Human Multidrug-Resistant Pathogen. Emerg Infect Dis, 30(4), 701–710. 10.3201/eid3004.231031

33. Meng, E. C., Goddard, T. D., Pettersen, E. F., Couch, G. S., Pearson, Z. J., Morris, J. H., & Ferrin, T. E. (2023). UCSF ChimeraX: Tools for structure building and analysis. Protein Sci, 32(11), e4792. 10.1002/pro.4792

34. Nuccio, S. P., & Baumler, A. J. (2007). Evolution of the chaperone/usher assembly pathway: fimbrial classification goes Greek. Microbiol Mol Biol Rev, 71(4), 551–575. https://doi.org/71/4/551 [pii] 10.1128/MMBR.00014-07

35. Ruhe, Z. C., Subramanian, P., Song, K., Nguyen, J. Y., Stevens, T. A., Low, D. A., Jensen, G. J., & Hayes, C. S. (2018). Programmed Secretion Arrest and Receptor-Triggered Toxin Export during Antibacterial Contact-Dependent Growth Inhibition. Cell, 175(4), 921–933 e914. 10.1016/j.cell.2018.10.033

36. Sachdeva, G., Kumar, K., Jain, P., & Ramachandran, S. (2005). SPAAN: a software program for prediction of adhesins and adhesin-like proteins using neural networks. Bioinformatics, 21(4), 483–491. 10.1093/bioinformatics/bti028

37. Sauer, K., Stoodley, P., Goeres, D. M., Hall-Stoodley, L., Burmolle, M., Stewart, P. S., & Bjarnsholt, T. (2022). The biofilm life cycle: expanding the conceptual model of biofilm formation. Nat Rev Microbiol, 20(10), 608–620. 10.1038/s41579-022-00767-0

38. Sia, C. M., Pearson, J. S., Howden, B. P., Williamson, D. A., & Ingle, D. J. (2025). Salmonella pathogenicity islands in the genomic era. Trends Microbiol, 33(7), 752–764. 10.1016/j.tim.2025.02.007

39. St-Pierre, F., Cui, L., Priest, D. G., Endy, D., Dodd, I. B., & Shearwin, K. E. (2013). One-step cloning and chromosomal integration of DNA. ACS Synth Biol, 2(9), 537–541. 10.1021/sb400021j

40. Wagner, C., & Hensel, M. (2011). Adhesive mechanisms of *Salmonella enterica*. Adv Exp Med Biol, 715, 17–34. 10.1007/978-94-007-0940-9_2

41. Wagner, C., Polke, M., Gerlach, R. G., Linke, D., Stierhof, Y. D., Schwarz, H., & Hensel, M. (2011). Functional dissection of SiiE, a giant non-fimbrial adhesin of *Salmonella enterica*. Cell. Microbiol., 13(8), 1286–1301. 10.1111/j.1462-5822.2011.01621.x

42. Zimmermann, L., Stephens, A., Nam, S. Z., Rau, D., Kubler, J., Lozajic, M., Gabler, F., Soding, J., Lupas, A. N., & Alva, V. (2018). A Completely Reimplemented MPI Bioinformatics Toolkit with a New HHpred Server at its Core. J Mol Biol, 430(15), 2237–2243. 10.1016/j.jmb.2017.12.007

