## Supplementary material for "Novel pESI-encoded autotransporter adhesin PeaP of epidemic *Salmonella* strains mediates adhesion, atypical biofilm formation, and poultry colonization": Suppl. Material

#### Supplementary Materials and Methods

*Construction of mutant strains and plasmids*

Gene deletion of *csgD* and *peaP* in SIN119944 was conducted by λ Red recombinase using a 3-step PCR method (Serra-Moreno et al., 2006). Genomic DNA of SIN 119944 and plasmid pKD3 together with oligonucleotides listed in Table 2 were used as templates. An amplimer was produced by annealing the 3 fragments by a 2-step PCR. Resulting fragments were further purified using the Monarch PCR purification kit. SIN 119944 [pKD46] was culture in LB with 20 mM L-arabinose for induction of λ Red recombinase, competent were prepared, and PCR products were electroporate, followed by selection on LB Cm agar. Loss of helper plasmid pKD46 was enforced by incubation at 42 °C and confirmed by sensitivity of clones to carbenicillin. Correct insertion of resistance cassettes and resulting deletion of genes of interest were checked by a colony PCR using the oligonucleotides listed in Table 2. Removal of resistance cassette for SIN 119944 *peaP*::CAT was conducted by transformation of pE-FLP in electro-competent cells of the respective strain and growth at 30 °C on LB agar plates containing 50 µg/ml carbenicillin. The method used here is described in detail in Serra-Moreno et al. (2006). Gene deletion of flagellar regulator *flhDC* was performed using an already existing mutant strain of STM (MvP2931) and oligonucleotides as listed in **Table S 1**. Large homologous regions for insertion of the *aph* resistance cassette and the respective specific gene region were amplified via Q5 Polymerase PCR and further purified using the Monarch PCR purification kit. The PCR product was transformed in electrocompetent cells of SIN 119944 [pKD46] induced by 20 mM L-arabinose for λ Red recombinase expression. Selection of successful insertion of resistance cassettes was performed by plating bacteria on LB agar plates containing 12.5 µg/ml chloramphenicol or 50 µg/ml kanamycin, respectively, and incubation at 37 °C. Loss of pKD46 was confirmed by incubation at 42 °C and check for growth in presence of 50 µg/ml carbenicillin. Correct insertion of resistance cassettes and resulting deletion of genes of interest were checked by a colony PCR using the oligonucleotides listed in **Table S 3**.

*Protein extraction of whole cells from biofilm and analysis via mass spectrometry*

For the analysis of SIN119944 proteomes, bacteria were grown as described above for `Analysis of biofilm formation by crystal violet assay´ for 144 h at 30 °C. After incubation, supernatant was collected and bacteria from supernatant were pelleted and recovered as planktonic fraction. Wells were washed three times with $H_2O_{dd}$ to remove residual planktonic bacteria and 150 µl PBS was added to the wells. Biofilm was scratched from the bottom of each well using plastic tips and resuspended in PBS. Biofilm mass of 96 wells was pooled, bacteria pelleted for 2 min at 8,000 x g and recovered as sessile fraction. Bacterial pellets were stored at -80 °C until further processing for proteomic analysis. Further sample preparation for proteomic analysis was done using the PreOmics iST Kit (PreOmics #P.O.00027, Planegg, Germany) according to the manufacturer’s instructions. For each pellet, 50 µl lysis buffer was used for lysis at 95 °C for 60 min. Subsequently, peptides were recovered in 2 x 25 µl elution buffer. Dried pellets were finally resolved in 100 µl LC load buffer, further diluted 1:10 in LC load buffer, transferred to HPLC vials, and subjected to perform reversed-phase chromatography on a Thermo Ultimate 3000 RSLCnano system connected to a TimsTOF HT mass spectrometer

(Bruker Corporation, Bremen) through a Captive Spray Ion source with approximately 100 ng. Peptides were separated on a Aurora Gen3 C18 column (25 cm x 75 µm x 1.6 µm) with CSI emitter (Ionoptics, Collingwood, Australia) at temperature of 40 °C. Elution of peptides from the column was realized via a linear gradient of acetonitrile from 10-35% in 0.1% formic acid for 44 min at a constant flow rate of 300 nl/min following a 7 min increase to 50%, and finally, 4 min to reach 85% buffer B. Eluted peptides were directly electro sprayed into the mass spectrometer at a electrospray voltage of 1.5 kV and 3 l/min Dry Gas. The MS settings of the TimsTOF were adjusted to positive Ion polarity with a MS range from 100 to 1700 m/z. The scan mode was set to DDA-PASEF. The ion mobility was ramped from 0.7 $Vs/cm^2$ to 1.5 in 100 ms. The accumulation time was also set to 100 ms. 10 PASEF ramps per cycle resulted in a duty cycle time of 1.17s. The target intensity was adjusted to 14,000, the intensity threshold to 1,200. The dynamic exclusion time was set to 0.4 min to avoid repeated scanning of the precursor ions, their charge state was limited from 0 to 5. The data were loaded to PeaksOnline Version 1.9 and analysed with the PeaksOnline workflow (DeNovo and DB Search) with Precursor mass error tolerance of 15 ppm, Fragment Mass error tolerance of 0.5 Da, CSS Error tolerance of 0.05 and a Missed Cleavage of 2. As modifications, Carbamidomethylation(C) (fix) and Oxidation(M) (variable) were chosen. Custom protein database 'SIN_old' based on the genome sequence of SIN 119944 was used for proteomic analyses. The results were filtered for peptides with an FDR of 1%, for proteins by the -10lgP of 20.

*Generation of antiserum against PeaP*

For generation of PeaP as antigen for immunization, *E. coli* C43 harboring p6443 was used for protein production. A culture in 2TY medium was grown with aeration at 37 °C to $OD_{600}$ of ca. 0.8. Expression was induced by addition of AHT to 10 ng/ml final concentration. Incubation was continued for 1.5 h and cell were harvested by centrifugation. The bacterial pellet was resuspended in 50 ml GST PBS supplemented with 5% glycerol (final), protease inhibitor cocktail, DNase and Lysozyme. After incubation for 30 min on ice, cells were lysed by ultrasonication using a Sonifier (Branson). Unbroken cells and cell debris were removed by centrifugation at 10,000 x g for 30 min. at 4 °C. The supernatant was then subjected to ultracentrifugation in a TI50.2 rotor at 50,000 rpm for 1 h at 4 °C. Supernatant and membrane pellet were further analysed SDS-PAGE to identify fractions with highest amounts of PeaP, and the membrane fraction was selected for further use. Proteins were separated by SDS-PAGE on 6% acrylamide gels using the Laemmli buffer system and subjected to staining by Coomassie brilliant blue. Due to the high molecular weight of PeaP and the absence of other protein in this size range, direct preparation was possible. Protein bands in the 180 kDa range were excised, and protein was recovered using an electro-eluter (BioRad Model 422) according to manufacturer's instructions. After dialyses to reduce SDS concentration, the eluted proteins were mixed with complete Freund's adjuvant for used as antigen for immunization of a New Zealand White rabbit using the 28 d SuperFast protocol (Davids Biotechnologie GmbH, Regensburg, Germany). The resulting immune serum was used directs in analyses as shown below.

*Immunostaining*

For immunolabeling, antibodies were used as follows: mouse anti *Salmonella* O:7 (Sifin, TR1305), mouse anti *Salmonella* H:r (Sifin, TR1416), rabbit anti PeaP (see above) diluted 1:500. As secondary antibodies, goat anti mouse-AlexaFluor488, goat anti mouse-AlexaFluor568, goat anti rabbit-AlexaFluor488, goat anti rabbit-AlexaFluor488, goat anti rabbit-AlexaFluor568 (all Jackson Laboratory) were used diluted 1:1,000.

For immunostaining of PeaP and flagella, cultures of respective strains were grown in CFA o/n at 37 °C, then adjusted to $OD_{600}$ of 0.01, fixed with 3% PFA in PBS for 15 min. Bacterial suspensions were spotted onto coverslips pretreated with 0.1% poly-L-lysine, coverslips were placed in wells of 24-well plates, and centrifuged at 500 x g for 5 min. Coverslips were washed and trice and incubated in blocking solution (2% goat serum in PBS) for 30 min. incubation with primary and secondary antibodies diluted in blocking solution was performed, with washing with PBS between incubations. Finally, coverslips were mounted with Fluoroshield, and imaged by widefield microscopy using AxioObserver system (Zeiss) with a 100x oil objective. For immunostaining of bacteria in biofilms, cultivation in CFA medium was performed directly on coverslips, that were fixed, processed as above, and imaged using a 40x air objective.

*Flow cytometry for analyses of PeaP surface expression*

Bacterial strains were cultured as before, and after sub-cultivation, 100 µl of the subculture was fixed with 100 µl of 3% PFA for 15 min at RT. Subsequently, blocking was performed for 10 min at RT with 200 µl of 100 mM ammonium sulfate to obtain a final concentration of 50 mM. Finally, 1 ml of PBS was added to the samples and measured in 1.5 ml reaction tubes. For analysis of PeaP surface expression, cultures were also prepared as An OD600 = 0.1 of the culture was pelleted for 2 min at 8.000 g and fixed in 200 µl 3% PFA/PBS for 20 min at RT. Samples were again pelleted and blocked with 200 µl blocking solution (HEPES, 0.2% Glycine, 0.1% BSA) for 30 min at RT. Antiserum rabbit αPeaP was added directly to blocking solution at 1:1,000 and incubated o/n in the orbital shaker at 4 °C. Next, samples were washed three times in PBS and incubated with goat α rabbit-Alexa488 at 1:2,000 dilution in blocking solution. After 1 h incubation in the dark the samples were again washed three times with 1 x PBS and resuspended in 400 µl PBS.

Flow cytometry was performed using the Attune NxT 4.2 system (ThermoFisher) with following settings: filter specifications: FSC/SSC 340/420, BL1 488/10 and YL2 620/15; voltages: FSC/SSC 340/420, BL1 400 andYL2 400; and a threshold: FSC 0.1x1000 and SSC 3.1x1000. The bacteria were detected at a flow rate of 12.5 µl/ml with 50,000 events for the 'bacteria' gate. The results of the flow cytometry were normalized using equation:

$n \text{ Alexa488-positive bacteria} * X\text{−median Alexa488} / n \text{ bacteria}.$

#### Suppl. Tables

**Table S 1.** Oligonucleotides used in this study

| designation | sequence (5´ - 3´) | application |
|---|---|---|
| Cm-5 | TGTGTAGGCTGGAGCTGCTTC | deletion of Δ*csgD* and *peaP*; check-PCR |
| Cm-3 | CATATGAATATCCTCCTTAG | deletion of Δ*csgD* and *peaP* |
| csgD-for | AAAAAGGTCTGGAAAATAAC | deletion of Δ*csgD,* check PCR |
| Cm-5-csgD-rev | CTAAGGAGGATATTCATATG TAAGGCCATGAAACGCTATC | deletion of *csgD* |
| Cm3-csgD-for | GAAGCAGCTCCAGCCTACACA ATGATGAAACTCCACTTTTT | deletion of *csgD* |
| SIN-GVI52_23260-for | TGGAAAACATTATTGCTCCT | deletion of *peaP* |
| Cm-5-peaP-rev | CTAAGGAGGATATTCATATGCATA ATAAGTCACTCAGTTT | deletion of *peaP* |
| SIN-GVI52_23270-rev | TCTGGTGTCAGAATTTCCTT | deletion of *peaP* |
| Cm-3-peaP-for | GAAGCAGCTCCAGCCTACACATGG AATTCGTTACAGTTTCT | deletion of *peaP* |
| SIN-GVI52_23260-for | TGGAAAACATTATTGCTCCT | check PCR for Δ*peaP* |
| SIN-GVI52_23270-rev | TCTGGTGTCAGAATTTCCTT | check PCR for Δ*peaP* |
| flhCD-delcheck-rev | CGTGATTTGGCCATCAGGCG | check PCR for Δ*flhDC* |
| SPA-flhCD-delcheck-for | ATCTATTATCCTGGCGTTAT | check PCR for Δ*flhDC* |
| k1-red-del | CAGTCATAGCCGAATAGCCT | check PCR, universal |

| | | |
|---|---|---|
| peaP SDM Del446 For | ACCAAAATTAGCAATAGTAAAC | SDM for p6756 |
| peaP SDM Del27 Rev | ATCAACAGACAATGCTG | SDM for p6756 |
| peaP SDM Del1478 For | CTCATAGTCAGGCAAGCAAG | SDM for p6757 |
| peaP SDM Del1383 Rev | ATCCAGATATGCGGTTTG | SDM for p6757 |
| peaP SDM Del1382 For | AGCTCTGGAAATGTGGTG | SDM for p6758 |
| peaP SDM Del447 Rev | TGCATCAACGACGCCATC | SDM for p6758 |
| peaP SDM Del955 For | CTCCTGACAGTGAACAGC | SDM for p6759 |
| peaP SDM Del632 Rev | GGTCGTGGAACTGATATTC | SDM for p6759 |

**Table S 2.** Design of chicken infection experiments

| group | infection | strain | n birds | n infected + n sentinel birds |
|---|---|---|---|---|
| 1 | yes | SIN 119944 | 25 | 15 + 10 |
| 2 | yes | SIN D*peaP* | 25 | 15 + 10 |
| 3 | yes | SIN 119944 [pGEN-*lux*] | 25 | 15 + 10 |
| 4 | yes | SIN Δ*peaP* [pGEN-*lux*] | 25 | 15 + 10 |
| 5 | no | - | 15 | - |

**Table S 3.** SIN detection in organs of infected chicken

| | SIN 119944 | SIN Δ*peaP* |
|---|---|---|
| Chicken infected (n) | 50 | 49 |
| SIN in caecum (%) | 100 | 100 |
| SIN in gallbladder (%) | 34 | 18.37 |

| | | |
|---|---|---|
| SIN in spleen (%) | 50 | 28.57 |
| SIN in liver (%) | 36 | 28.57 |

#### Suppl. Figures and Legends

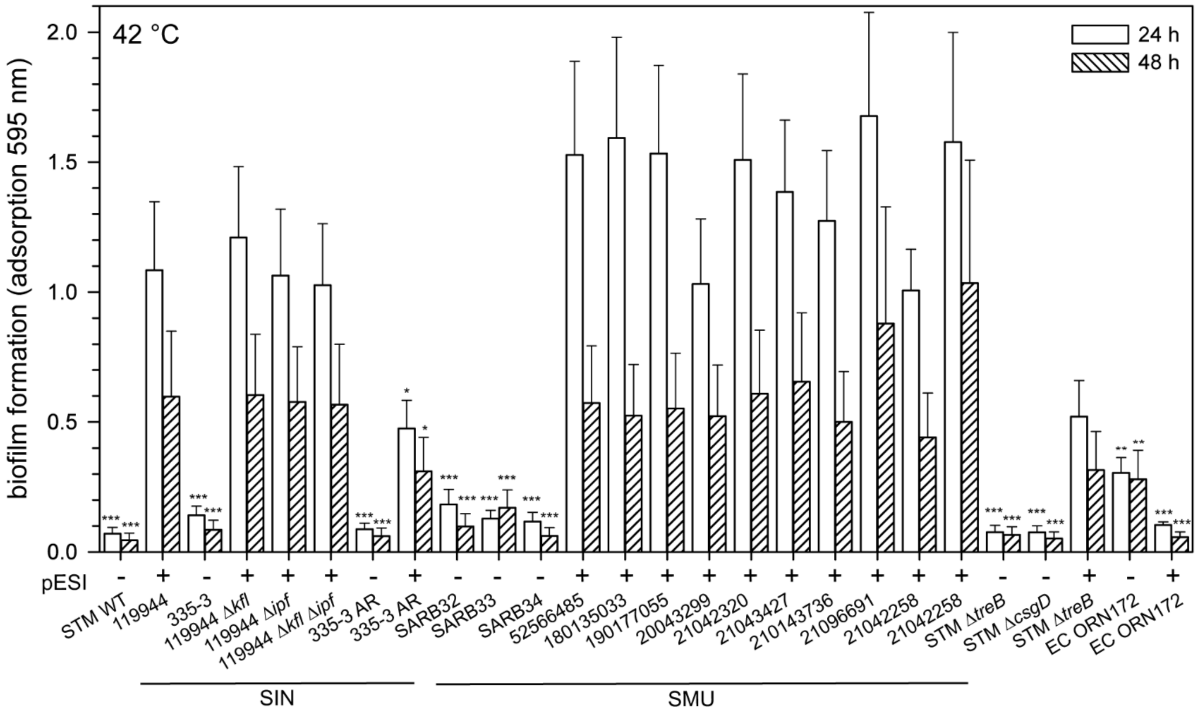


**Figure S 1. Presence of pESI mediates atypical biofilm formation.** Various *S. enterica* isolates of serovars Typhimurium (STM), Infantis (SIN), Muenchen (SMU), and *E. coli* (EC) ORN172 and isogenic mutant strains were analysed for biofilm formation at 42 °C. The absence or presence of pESI is indicated by – and +, respectively. Biofilm formation was quantified as described for **Figure 1**. Means and standard deviations of at least three biological replicates are shown. Statistical analyses for comparison to SIN 119944 were performed by ANOVA and results are indicated as: *, $p < 0.05$; **, $p < 0.01$; ***, $p < 0.001$, all other data are not significantly different (not indicated).

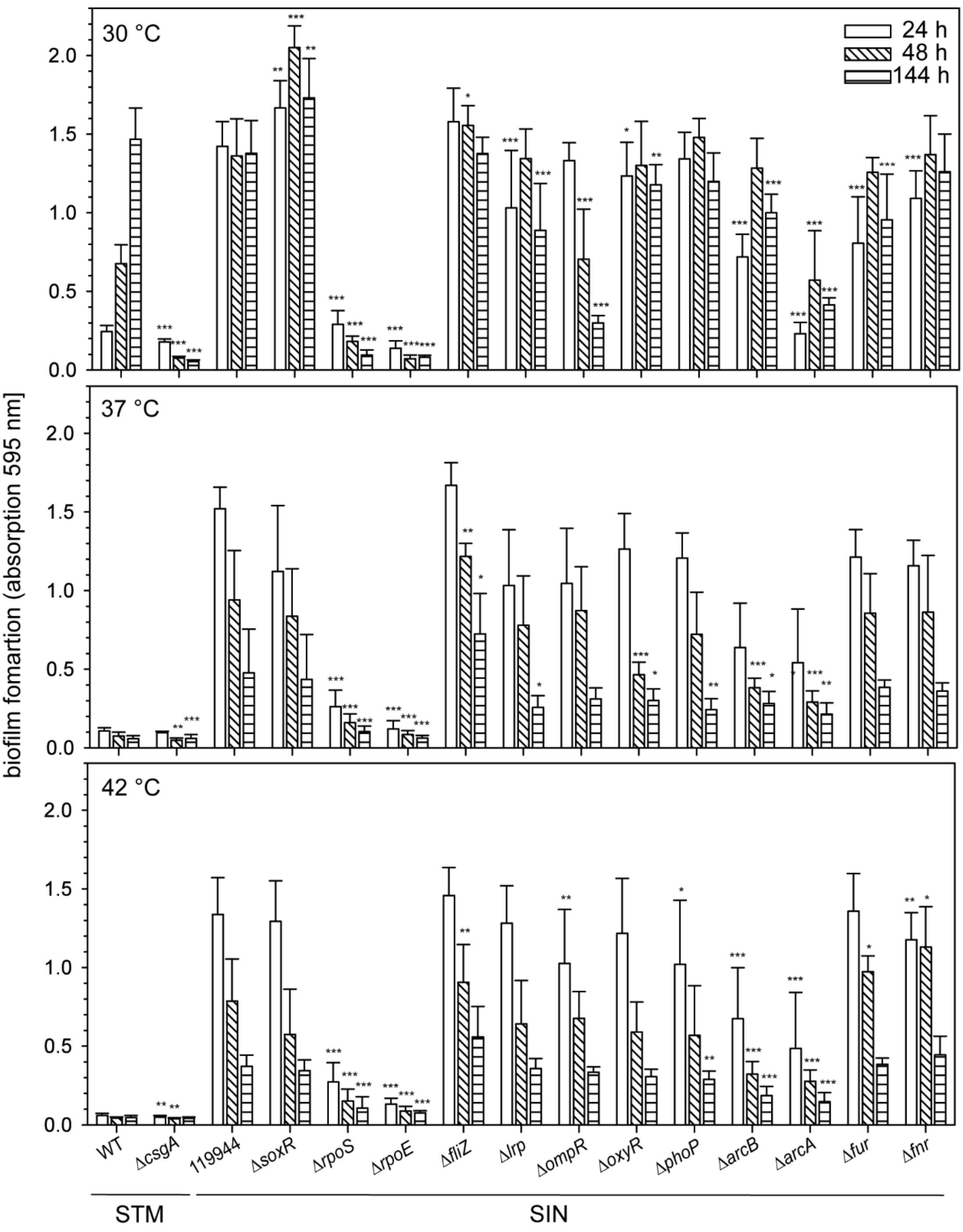


**Figure S 2. Role of global regulatory systems in atypical biofilm formation of SIN 119944.** Biofilm formation of STM WT, STM Δ*csgD*, SIN 119944, and various mutant strains isogenic to SIN 119944 as indicated was analysed as described for **Figure 1**. Means and standard deviations of at least three biological replicates are shown. Statistical analyses for comparison to STM WT or SIN 119944 were performed by ANOVA and results are indicated as: *, $p < 0.05$; **, $p < 0.01$; ***, $p < 0.001$, all other data are not significantly different (not indicated).

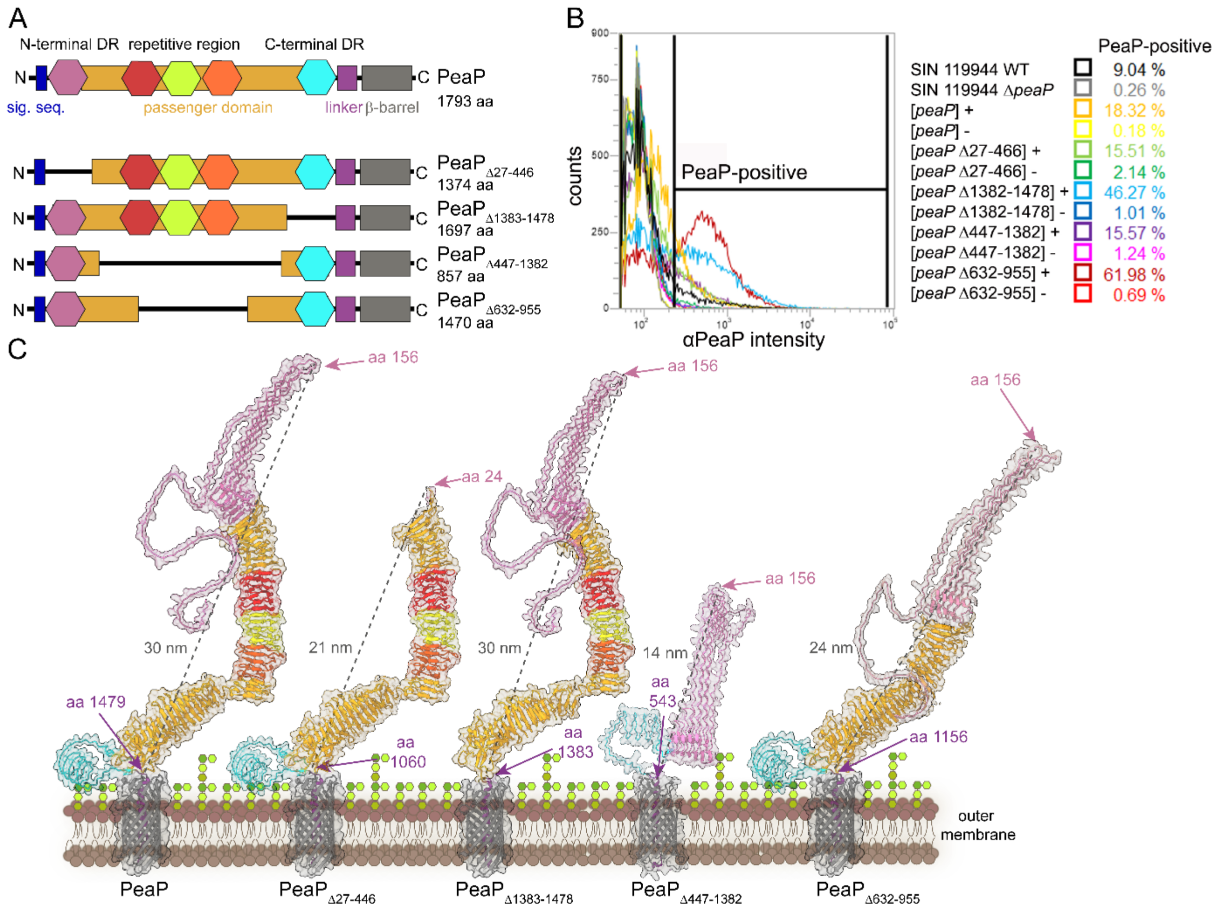


**Figure S 3. Deletion of domains of PeaP and surface expression of PeaP alleles**. **A**) Domain organization of PeaP and position of domain deletions analysed in this study. Deletions in PeaP comprised aa 27-466 (N-terminal disordered region), aa 1382-1478 (C-terminal disordered region), aa 447-1382 (passenger domain), aa 632-955 (passenger domain). **B**) Flow cytometry analyses of surface expression of WT PeaP and various mutant alleles. Surface expression was analyzed in SIN 119944 Δ*peaP* complemented with various plasmids as indicated. The tet-on expression system was used for episomal expression of WT *peaP* and mutant alleles of *peaP* with internal deletions as indicated. Strains were subcultured with (+) or without (-) addition of AHT, fixed, subjected to labelling and with antibodies against PeaP, and analysed by flow cytometry to quantify the percentage of PeaP-positive bacteria. **C**) The structures of PeaP and various deletion alleles were predicted using AlphaFold 3. The calculated distance between the α-helical linker (aa1479 in WT PeaP) and distal part of PeaP is indicated in nm.

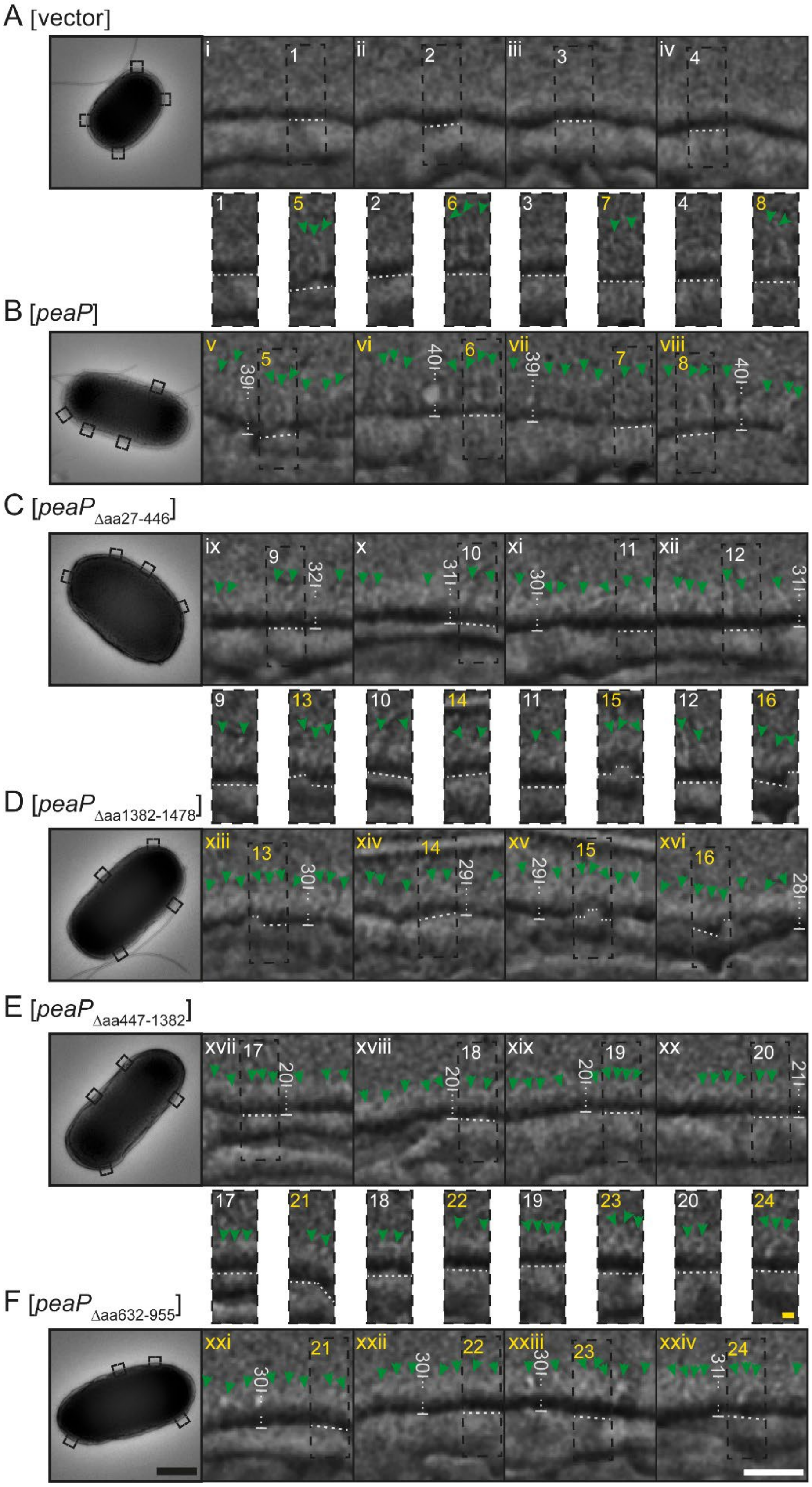


**Figure S 4. Surface expression of truncated forms of PeaP**. SIN 119944 Δ*peaP* harboring the empty vector [vector] (**A**), or plasmids for AHT-inducible expression of WT *peaP* [*peaP*]

(**B**) or deletion alleles of *peaP* as indicated (**C, D, E, F**) were used. Strains were cultured as described for **Figure 6** and $P_{tetA}$-controlled expression of was induced by addition of 1 µg/ml AHT after subculture for 1 h. Negative stain TEM of various strains was performed as for **Figure 6**. Green arrowheads indicate representative PeaP molecules. Scale bars: overview images (black bar), 500 nm; enlarged images (white bar), 50 nm; detail images (yellow bar), 10 nm.

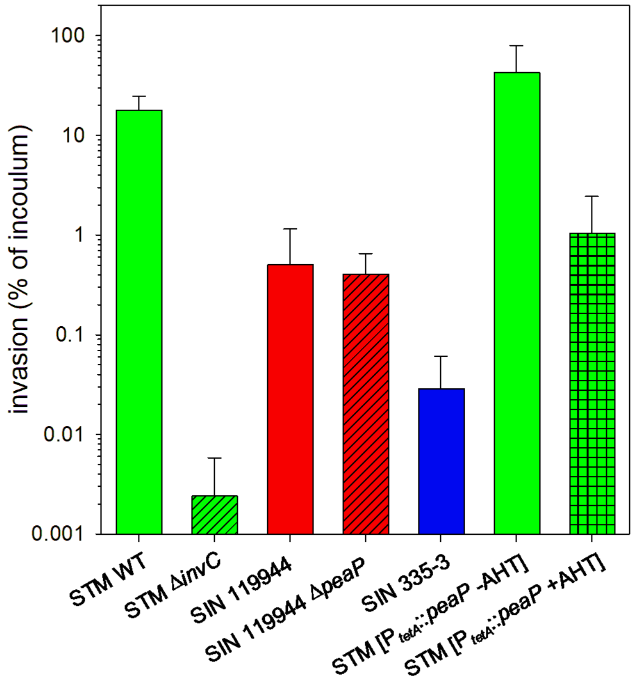


**Figure S 5. PeaP is dispensable for invasion of polarized epithelial cells.** The polarized epithelial cell line MDCK was used as host cells for infection by STM or SIN strains as indicated. STM harboring a plasmid for synthetic expression of *peaP* was used after subculture without or with addition of AHT for expression of *peaP* under control of $P_{tetA}$. After infection for 60 min, cells were washed and medium containing 100 µg/ml Gentamicin was added to kill non-internalized bacteria. Infected MDCK cells were lysed and lysates plated for CFU determination. Invasion is expressed as percentage of internalized inoculum, and means and standard deviations of three replicates are shown.

### Suppl. References

Serra-Moreno, R., Acosta, S., Hernalsteens, J. P., Jofre, J., & Muniesa, M. (2006). Use of the lambda Red recombinase system to produce recombinant prophages carrying antibiotic resistance genes. *BMC molecular biology*, *7*, 31. https://doi.org/10.1186/1471-2199-7-31
